# Artemisia Database: A Comprehensive Resource for Gene Expression and Functional Insights in *Artemisia annua*

**DOI:** 10.1101/2025.05.21.655314

**Authors:** Ayat Taheri, Fabricio Almeida-Silva, Yaojie Zhang, Xueqing Fu, Ling Li, Yuliang Wang, Kexuan Tang

## Abstract

*Artemisia annua* is renowned for producing artemisinin, a compound that revolutionized malaria treatment and holds therapeutic promise for other diseases, including cancer and diabetes. However, low natural yields of artemisinin remain a major bottleneck, necessitating a deeper understanding of the genetic and regulatory networks involved in its biosynthesis. Although several transcriptomic studies on *A. annua* exist, they are often limited in scope, and a comprehensive, tissue-resolved gene expression resource has been lacking. Here, we present the Artemisia Database (Artemisia-DB)—a high-resolution expression atlas constructed from an extensive integration of publicly available RNA-seq datasets. The database provides transcript- and gene-level abundance estimates across major tissues and includes functional annotations such as Gene Ontology (GO) terms, KEGG pathways, and InterPro domains. As a case study, we investigated the coexpression profile of HMGR (3-hydroxy-3-methylglutaryl-CoA reductase), a key enzyme in the mevalonate pathway and an early step in artemisinin biosynthesis. Coexpression analysis in leaf tissue revealed a subset of Auxin Response Factor (ARF) transcription factors strongly correlated to HMGR. This finding suggests a potential regulatory link between auxin signaling and artemisinin biosynthesis, providing new hypotheses for functional validation. Artemisia-DB is freely accessible at https://artemisia-db.com and offers an interactive interface for exploring expression data, functional annotations, transcription factors, CRISPR targets, and more. By combining high-quality transcriptome data with regulatory and functional insights, Artemisia-DB serves as a valuable resource for the plant research community and facilitates deeper investigation into the transcriptional dynamics and specialized metabolism of *A. annua*.

## Introduction

*Artemisia annua* L., commonly known as sweet wormwood, has had a profound impact on global health through its production of artemisinin, a compound that revolutionized malaria treatment [1]. Beyond its well-established role in combating malaria, artemisinin and its derivatives have demonstrated potential in treating a variety of other diseases, including cancer and diabetes, highlighting the broad therapeutic potential of this compound [2]. However, the natural production of artemisinin in *A. annua* is rather low, posing a substantial barrier to fulfilling worldwide demand and encouraging research efforts to enhance its production [2]. Achieving this goal requires a deeper understanding of the genetic mechanisms that regulate artemisinin biosynthesis.

Despite extensive research on the medicinal properties of *A. annua*, the gene expression patterns across its various tissues have been examined in only a limited number of studies, often relying on small or narrowly focused datasets. While individual experiments are typically designed to test specific hypotheses, integrating diverse transcriptomic datasets has the potential to reveal novel regulatory patterns and biological insights that may not be evident from any single study. The development of comprehensive gene expression databases is therefore crucial to bridging this knowledge gap and facilitating future discoveries [3]. While previous efforts to create expression databases for plant species (e.g., PlantExp and IMP [4], [5]) have been instrumental, they either exclude *A. annua* or rely on outdated genomic data with limited RNA-seq samples, leaving a significant gap in our understanding of its transcriptome.

This study fills such a gap by first enhancing the quality of the transcriptome assembly through the integration of diverse RNA-seq datasets, followed by a unique, in-depth analysis to identify tissue-specific genes, transcription factors (TFs), and other key regulatory elements. The result is a comprehensive expression atlas for *A. annua*, providing valuable insights into global gene expression patterns, functional annotations—including GO terms, KEGG pathways, and InterPro domains—and the characterization of tissue-specific TFs that govern developmental and metabolic processes.

To further support the research community, we have developed a dedicated online platform, Artemisia Database (Artemisia-DB) (https://artemisia-db.com), offering detailed transcript- and gene-level abundance estimates. This resource enables researchers to explore differential transcript usage across key tissues, including leaves, flowers, and stems. The database includes both raw counts and normalized expression values, facilitating in-depth comparative analyses. Moreover, it provides a curated collection of genes involved in artemisinin biosynthesis, transcriptional regulation, and epigenetic modifications. Users can also explore tissue-specific expression patterns of TFs identified from databases such as PlantTFDB and Pfam.

By integrating high-quality transcriptome data with functional annotation and regulatory insights, Artemisia-DB represents an unparalleled resource for studying *A. annua*’s transcriptional dynamics and specialized metabolism, offering a powerful tool for researchers worldwide.

## Results and discussion

### Data Summary and Quality Metrics

The multi-step computational workflow developed to construct Artemisia-DB is shown in Figure 1. In total, 267 RNA-Seq datasets from 23 different projects and four available Single Molecule, Real-Time (SMRT) sequencing samples were downloaded, all representing various ecotypes of *A. annua*. Most of the data (134 out of 271 samples) came from the “Huhao1” ecotype, with information missing for 60 entries. Leaf samples were predominant (194), followed by root (40), flower (20), and stem (14), while only three samples were available for petiole. Samples for bud and seed having only one replication (two samples) were excluded from further study. Trichomes and epidermis samples (one each) were classified as leaf. The library layout for 237 datasets was paired-end, with the remaining 30 being single-end. The samples spanned diverse developmental stages (from one month to one year) and under various conditions (e.g., Jasmonic acid, light, high temperature, graphene, and UV treatment).

**Figure 1.**
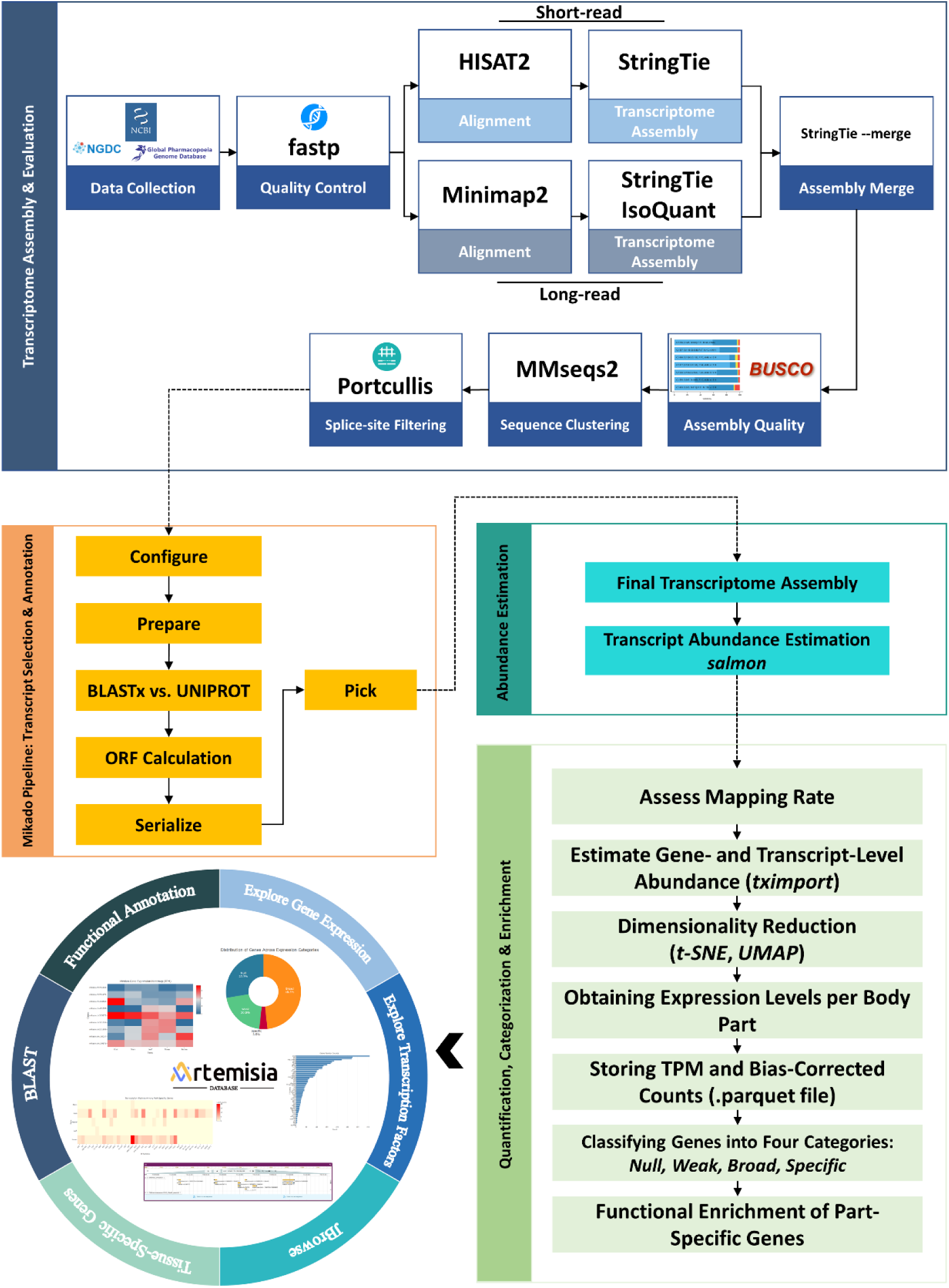
Workflow of the Artemisia Database development and analysis

RNA-Seq data quality was ensured by applying minimum thresholds of a mean read length greater than 40 bp and a Q20 rate above 80%, which were met by all 266 samples (Figure 2A). A scatter plot with marginal violin plots (Figure 2A) shows a tight Q20 rate distribution peaking at 97.5–99.5%, confirming high sequencing quality, and a bimodal total bases distribution with peaks at ∼1000 million (lower-depth) and 5000–7,500 million (higher-depth), slightly skewed right due to a few high-depth samples (up to 12,500 million). The scatter plot reveals a dense cluster at Q20 rates of 97.5–99.5% and total bases of 5000–7,500 million, with a smaller cluster of lower-depth samples. This depth heterogeneity impacts detection sensitivity—insufficient depth reduces power for identifying differential expression of low-abundance transcripts, while excessive depth yields diminishing returns [6], [7], [8]. Unfortunately, due to specific goals or other considerations of some experiments, a number of them lacked the minimum required replications and were consequently removed from this study. Additionally, the metadata uploaded for the samples was sometimes incomplete or incorrect, with errors such as specifying the wrong sequencing layout. To ensure that such data can be effectively used in future research, researchers must provide accurate and comprehensive metadata when submitting their datasets, as this information is essential for proper data interpretation and reproducibility. Overall, a dataset of 12.54 billion reads and 1.60 terabases (Tb) of clean data was obtained for further analysis.

**Figure 2.**
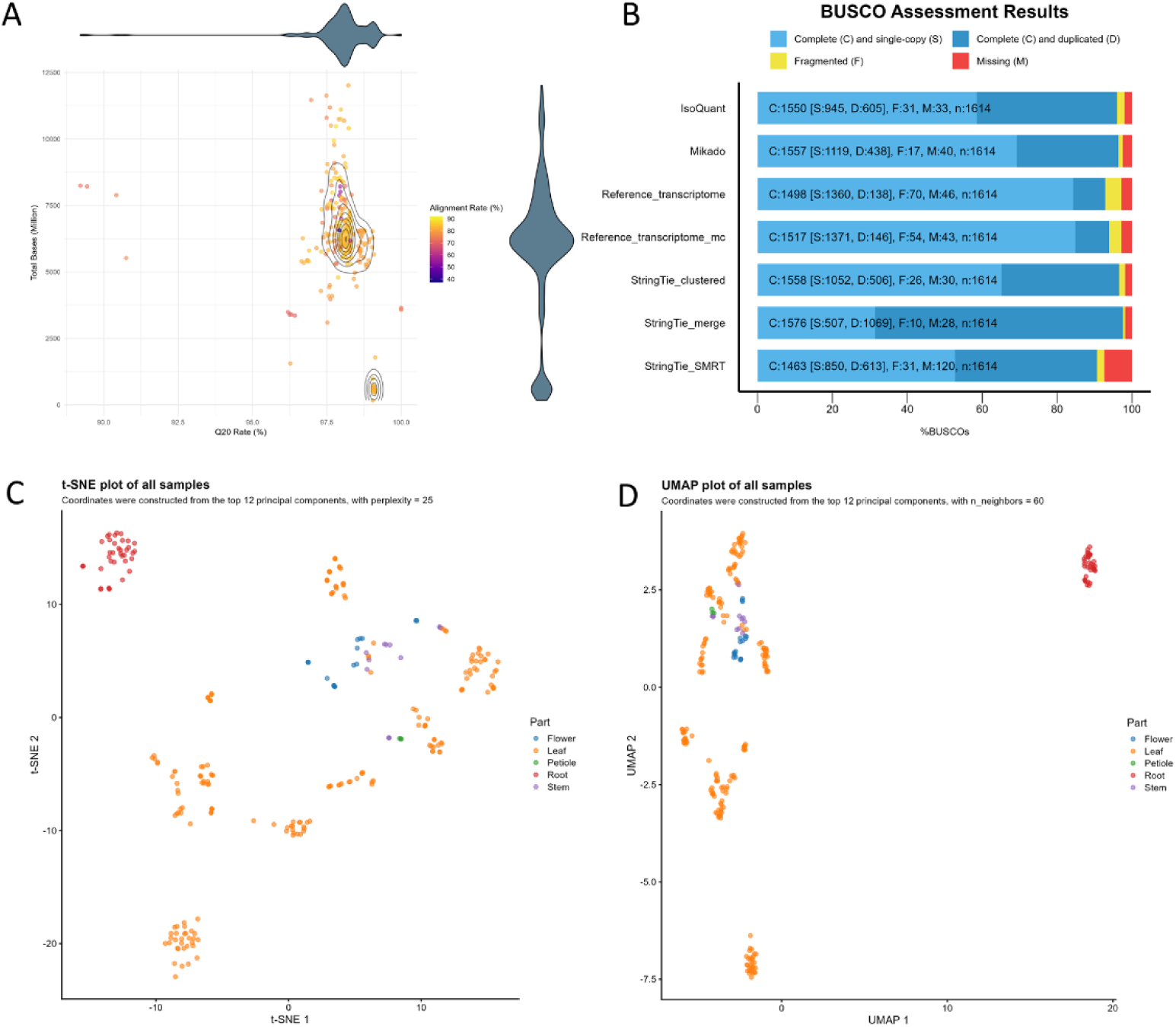
(A) Scatter plot of Q20 rate (%) vs. total bases (million) with alignment rates (%) as color, marginal violin plots, and density contours for 266 RNA-seq samples. (B) BUSCO summary assessing the completeness of all assemblies. (C) t-SNE plot of RNA-seq samples based on the top 12 principal components for dimensionality reduction. (D) UMAP plot of RNA-seq samples based on the top 12 principal components for dimensionality reduction.

### Alignment of RNA-Seq Reads to the *A. annua* Genome

HISAT2 was used to align the high-quality RNA-Seq data obtained in the previous step to the latest *A. annua* genome assembly [9], ensuring accurate read mapping and enabling comprehensive transcriptome analysis. Alignment rates across samples were high, with mean and median rates of 80.95% and 82.75%, respectively (Figure 2A)Figure 2. The alignment rates ranged from a minimum of 37.33% to a maximum of 92.08%, with an interquartile range (IQR) of 5.01%, highlighting that the middle 50% of the data had relatively consistent alignment rates. Notably, only one sample achieved an alignment rate above 90%, and similarly, just one sample had an alignment rate below 50%, which was excluded from further analysis to ensure the strength and reliability of the dataset.

### Transcriptome Assembly and Merging

To ensure a robust starting point, the quality of the reference transcriptome was initially evaluated using BUSCO, alongside another manually curated annotation [10]. Recognizing the potential of leveraging the extensive data available, a strategy was devised to construct a comprehensive transcriptome by merging annotations generated from various tools. This approach incorporated both short-read and long-read sequences to maximize coverage and accuracy. The BUSCO results for all assemblies are presented in Figure 2B.

Initially, StringTie was used to assemble individual transcriptomes for each short-read sample, followed by the “--merge” function to consolidate these into a final merged transcriptome [11]. For long-read sequences, two assemblers were employed: StringTie and IsoQuant. The quality and completeness of all transcriptome assemblies, as well as the reference transcriptome, were assessed using BUSCO. The StringTie-merged transcriptome exhibited superior completeness, identifying 97.6% of BUSCO genes as complete, compared to 92.9% for the reference transcriptome, 93.9% for the manually curated reference transcriptome, 90.7% for the StringTie long-read assembly, and 96.1% for the IsoQuant transcriptome (Figure 2B). However, the StringTie-merged transcriptome also displayed a higher proportion of duplicated BUSCO genes (66.2%), likely reflecting the capture of different isoforms by StringTie rather than true gene duplications [12]. Notably, the StringTie-merged transcriptome had the lowest percentages of fragmented (0.6%) and missing (1.8%) BUSCO genes.

MMseqs2 was employed to reduce the number of duplicated genes in the StringTie-merged assembly. This approach significantly reduced the duplication rate, lowering the number of complete and duplicated BUSCO genes from 1,069 in the StringTie-merged assembly to 506 — a reduction of approximately 52% after duplicate removal. To leverage all generated assemblies, the Mikado pipeline was used to merge them [13]. The resulting Mikado assembly exhibited a significantly greater number of complete BUSCO genes compared to most assemblies, except for the StringTie-merged assembly. However, Mikado’s advantage lay in its lower duplication rate (27.1% compared to 66.2% for StringTie). Consequently, the Mikado assembly was selected as the final assembly for downstream analyses.

### Clustering of RNA-Seq Samples Using t-SNE and UMAP

We used both t-SNE and UMAP dimensionality reduction techniques to explore the clustering of RNA-Seq samples based on their transcript abundance profiles, leveraging the top 12 principal components that explained over 70% of the total variance in the dataset (S1A). Based on visual inspection of the results, a perplexity of 25 was chosen for t-SNE, while the optimal number of nearest neighbors for UMAP was set to 60 (Figure S 1A, B). The t-SNE (Figure 2C) and UMAP (Figure 2D) plots both visually reveal distinct groupings of samples, suggestive of tissue-specific expression patterns. In the t-SNE plot, samples appear to form well-defined groups, while the UMAP plot displays similarly consistent patterns, often with more compact clustering. These visual trends indicate potential underlying transcriptomic differences between tissues.

Notably, root samples consistently appear in tightly grouped clusters across both methods, suggesting a strong and homogeneous expression profile. Petiole and stem samples also tend to cluster near one another, and flower samples often appear adjacent to stem clusters, possibly reflecting shared developmental pathways or biological similarity. These patterns align with observations in previous studies, such as in soybean, where nodule and root tissues, as well as embryo and endosperm, showed overlapping expression characteristics [3].

In contrast, leaf samples display a more dispersed pattern in both visualizations. This spread may reflect the dynamic and variable nature of leaf transcriptomes, which are sensitive to environmental factors such as light, temperature, and stress [14], [15]. Additional variation may stem from the large number of leaf samples in the dataset, differences in ecotypes and developmental stages, or treatment conditions.

It is important to note that both t-SNE and UMAP are nonlinear methods that emphasize local structure and neighbor preservation; as such, spatial distances and exact cluster boundaries in the 2D projections should be interpreted cautiously and complemented by other analytical approaches [16], [17].

### Global Gene-expression patterns and tissue-specific genes

To comprehensively analyze gene expression patterns and regulatory mechanisms across various tissues of *A. annua*, we classified all genes into four distinct categories: null, broad, weak, and specific. This classification was determined using the tissue specificity index (τ index) and median TPM values, as detailed in the methods section. The τ index, which ranges from 0 for housekeeping genes to 1 for tissue-specific genes, serves as a measure of tissue specificity in gene expression [18]. Originally developed to identify genes with midrange expression profiles—representing over 50% of all expression patterns—the τ index provides crucial insights into gene function. More recently, an extended τ approach has been proposed to enhance the identification of tissue-specific genes across multiple tissues [19].

The distribution of genes across the different expression categories is presented in Figure 3A. The broad category was the most prevalent, comprising 20,717 genes characterized by τ < 0.85 and median TPM > 5. These broadly expressed genes are likely involved in essential cellular processes, serving as part of the core transcriptome required for the maintenance and survival of *A. annua* across different tissues. The null expression category was the second largest, encompassing 11,889 genes. These genes exhibited low expression levels (median TPM < 1 in all tissues) under the studied conditions. In the weak expression category, 8,767 genes were identified, all with a median TPM < 5 in all tissues. These genes may play roles in specialized functions or fine-tune physiological processes under particular conditions. Finally, the specific expression category included 1,514 genes with τ ≥ 0.85 and median TPM > 5. These tissue-specific genes are of particular interest, as they likely contribute to the specialized functions and development of specific tissues within *A. annua*.

**Figure 3.**
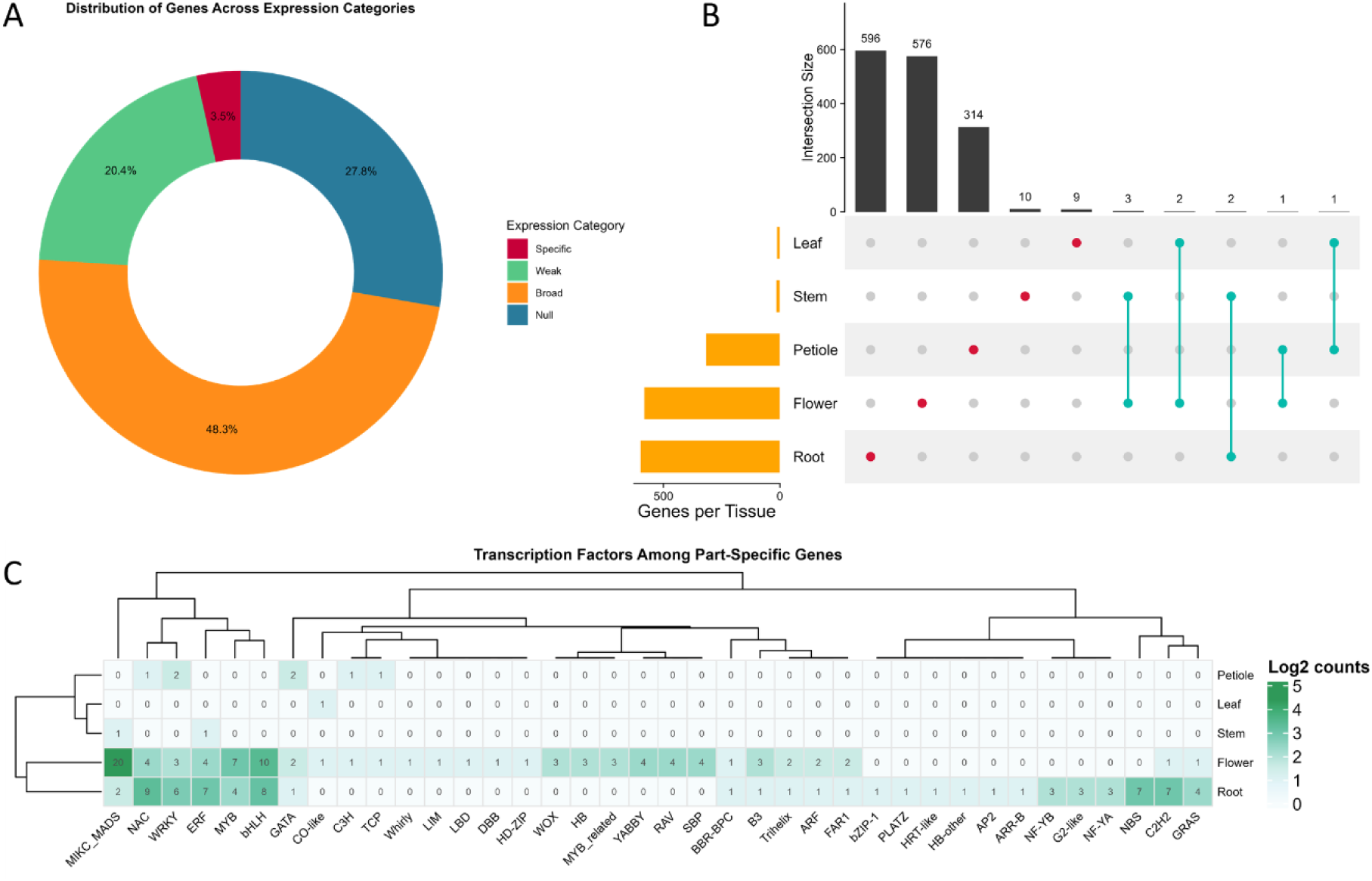
(A) Distribution of genes across four expression categories based on τ index and median TPM values: (1) Null expression (median TPM < 1 in all tissues); (2) Weak expression (median TPM < 5 in all tissues); (3) Broad expression (τ < 0.85, median TPM > 5); and (4) Body part-specific expression (τ ≥ 0.85, median TPM > 5). (B) Upset plot showing the overlap of tissue-specific gene expression across tissues. (C) Heatmap of transcription factor distribution among tissue-specific genes, identified using PlantTFDB and Pfam databases.

To pinpoint the tissues expressing these specific genes, we performed an analysis of gene expression patterns across different tissues (Figure S 1D). Our findings revealed that the root exhibited the largest number of tissue-specific genes, followed by the flower and petiole. In contrast, the stem and leaf had the smallest counts of tissue-specific genes. Notably, some genes were expressed in multiple tissues, with the flower and stem showing the greatest overlap, sharing three genes (Figure 3B).

The abundance of root-specific genes suggests significant specialization, likely reflecting the unique physiological functions of this tissue. Conversely, the minimal number of specific genes in the leaf and stem could indicate either a shared transcriptome with other tissues or less tissue-specific gene expression under the conditions studied.

### Functional Enrichment Analysis

We performed gene ontology (GO) and InterPro protein domain enrichment analyses to further investigate the functional roles of tissue-specific genes, using all expressed genes as the background set (Figure S 2). The results highlighted distinct functional categories associated with each tissue.

The analysis revealed that root tissues are enriched with terms such as "Heme binding", "Monooxygenase activity", and "Oxidoreductase activity". These activities are integral to metabolic processes and defense mechanisms within the root. The identification of "Cytochrome P450" and "Fe(2+) 2OG dioxygenase domain" further indicates the presence of active secondary metabolism in the root tissue. These findings align with existing research that highlights the diverse oxidoreductase activities in root tissues, including monooxygenases, dioxygenases, and peroxidases, which are crucial for plant metabolism and stress responses. The presence of heme-binding proteins, such as cytochrome P450, underscores their importance in oxidative reactions and stress tolerance, which are critical for the plant’s defense against environmental stressors [20]. These proteins, which rely on heme as a cofactor, play a central role in the transformation of superoxide anions within the plant’s antioxidant system. Indeed, the heme content of plants is directly related to their tolerance to environmental stressors [21].

Our analysis identified enriched terms such as "Retrotransposon gag domain" and "MULE transposase domain", indicating a significant involvement of transposable elements (TEs) in the petiole tissue. Although traditionally regarded as genomic parasites, TEs can be co-opted for beneficial functions in the host, such as regulating gene expression and maintaining genome stability. Long terminal repeat (LTR) retrotransposons, particularly the gag and pol genes, are crucial for retrotransposition, while Mutator-like elements (MULEs), have been repurposed in plants to form gene families like FAR1-related sequence (FRS) and MUSTANG (MUG), which are vital for development and environmental response, including light signal transduction, regulating the flowering time, and chloroplast division [22]. These findings suggest that TEs may play a critical role in regulating gene expression and responding to environmental stimuli.

The flower-specific genes were enriched in terms like "Transcription factor (K-box)" and "Transcription factor (MADS-box)", both crucial for flower development, morphology, and coloration [23], [24]. The presence of "Plant non-specific lipid-transfer protein" (nsLTP) points to roles in lipid metabolism and protein processing, vital for flower maturation. NsLTPs are essential for membrane stabilization and resistance to various stresses [25].

Enriched terms in the leaf tissue, such as "Transcription factor MYC/MYB N-terminal", further supports the leaf’s role in synthesizing secondary metabolites, which are important for plant defense and growth regulation. Numerous studies have highlighted the role of MYB transcription factors in regulating the artemisinin biosynthesis pathway. For example, AaMYB108 in *A. annua* is a key regulator of artemisinin production, interacting with AaGSW1 and modulating its function through interactions with AaCOP1 and AaJAZ8, thereby integrating light and jasmonic acid (JA) signals to regulate terpene biosynthesis [26]. Additionally, AaMYB1 significantly enhances artemisinin production by overexpressing essential biosynthetic genes and promoting trichome development. This transcription factor, along with its orthologue AtMYB61 in *Arabidopsis thaliana*, plays a crucial role in trichome initiation, root development, stomatal aperture, and gibberellin biosynthesis and degradation, making it a valuable target for genetic engineering to increase artemisinin yield [27].

### Characterization of Tissue-Specific Transcription Factors

Given their crucial roles in regulating gene expression across various tissues and the wealth of existing research emphasizing the importance of transcription factors (TFs) in *A. annua*, we conducted an in-depth analysis of TF distribution among tissue-specific genes. This analysis combined TFs identified through BLASTx against those provided by PlantTFDB (Figure S 3A) with those identified using Pfam IDs (Figure S 3B) from 63 different TF families (Figure 3C). The results revealed distinct tissue-specific expression patterns for several TF families, underscoring their roles in specialized metabolic pathways.

Notably, both root and flower tissues exhibited a significant presence of transcription factors across various families. In root tissue, these TFs spanned 24 different families, while in flower tissue, they were distributed across 29 families. In contrast, stem and leaf tissues presented the lowest number of tissue-specific TFs. Certain TF families were found to be exclusively expressed in particular tissues. For instance, the SBP, YABBY, RAV, and HD-ZIP families were specifically observed in flower tissue, likely reflecting their roles in flower development, hormone signaling transduction, and vegetative to reproductive phase transition [28], [29], [30], [31], [32]. Similarly, the HB-other, G2-like, NF-YA, and NF-YB families, exclusive to root tissue, may play crucial roles in root development and response to environmental stress [33], [34], [35].

Our analysis also highlighted the dominance of certain TFs within specific tissues. In root tissue, NAC, bHLH, ERF, NBS, and C2H2 were the most prominent, whereas in flower tissue, MIKC_MADS, bHLH, MYB, and SBP were dominant. The absence of ERF family TFs in the leaf, despite previous reports of their involvement in artemisinin biosynthesis and trichome development in *A. annua*, may be attributed to their broader functional roles across multiple tissues. This suggests that while ERF TFs are crucial for processes such as artemisinin biosynthesis, their activity is not confined to a single tissue, thereby not meeting the criteria for tissue specificity in our analysis.

In conclusion, our comprehensive analysis of transcription factor distribution across various tissues in *A. annua,* has unveiled a complex tapestry of gene regulation. The identification of distinct TF families dominant in specific tissues, alongside the broader functional roles of certain TFs like ERF, underscores the complexity of gene regulation in this species. These findings, made accessible through the Artemisia-DB, not only enhance our understanding of the molecular mechanisms underlying artemisinin biosynthesis and plant development but also provide a valuable resource for future research aimed at manipulating these pathways for improved artemisinin production.

### Case Study: Coexpression Analysis of HMGR Suggests Involvement of Auxin-Responsive ARF Transcription Factors in Leaf Tissue

To explore transcriptional regulators potentially modulating HMGR (3-hydroxy-3- methylglutaryl-CoA reductase)—a key enzyme in the mevalonate (MVA) pathway and a precursor step in artemisinin biosynthesis—we used the tools available in the Artemisia-DB (Figure 4). Using its BLAST module, we identified the HMGR gene from the reference genome based on its coding sequence.

**Figure 4.**
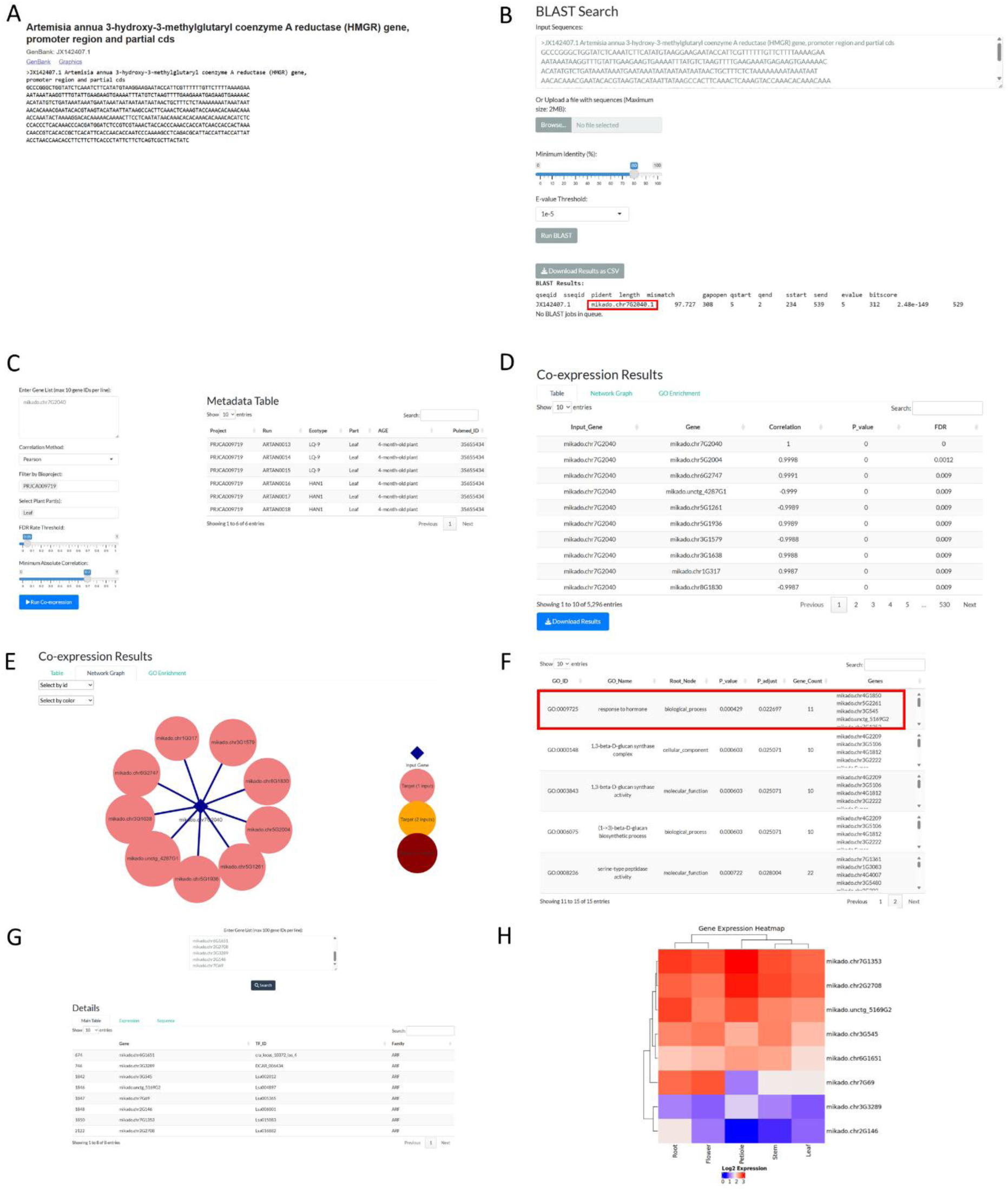
Case study of the *HMGR* gene in Artemisia-DB: (A) *HMGR* gene sequence from NCBI; (B) BLAST results against Artemisia-DB to identify the corresponding gene ID; (C) selection of co-expression analysis parameters (Pearson method, leaf samples, FDR ≤ 0.05, |correlation| ≥ 0.7); (D) co-expression results table; (E) co-expression network of the input gene and its top 10 co-expressed genes; (F) GO enrichment highlighting "response to hormone" and using the corresponding gene IDs; (G) TFDB search showing several ARF transcription factors; (H) heatmap of ARF TF expression across tissues.

Subsequently, we performed coexpression analysis using RNA-seq data specifically from leaf tissue. Among the top coexpressed genes, we filtered for those annotated with the Gene Ontology (GO) term “response to hormone” (GO:0009725), resulting in 11 genes. Cross-referencing these with the transcription factor database (TFDB) section of Artemisia-DB revealed that eight of them belong to the Auxin Response Factor (ARF) family.

To assess their broader transcriptional activity, we examined the tissue-specific expression patterns of these ARFs. Five of the eight showed consistently high median expression across major tissues. A dendrogram based on gene expression profiles indicated that leaf and stem tissues cluster closely, implying a shared regulatory environment where these ARFs may play a functional role.

These results align with previous findings that place HMGR at a regulatory intersection in artemisinin biosynthesis. It is reported that abscisic acid (ABA) treatment significantly upregulates HMGR expression and increases artemisinin content [36]. Likewise, studies by Ji et al. (2014) and Lu et al. (2013) demonstrated that HMGR expression can be enhanced through transcription factors such as AabHLH1 and AaORA, as well as through overexpression approaches [37], [38]. Although Guo et al. (2022) described a miR160–ARF1 module that enhances AaDBR2 expression and artemisinin accumulation, no direct connection has been established between ARF transcription factors and HMGR regulation.

Our analysis introduces new evidence suggesting that a subset of ARFs may be functionally linked to HMGR expression, at least at the transcriptional coexpression level in leaf tissue. While direct regulatory interactions remain to be validated, these results suggest that auxin-responsive ARFs could be part of a broader hormone-responsive regulatory network influencing the MVA pathway and artemisinin biosynthesis in *A. annua*.

This case study exemplifies the power of the Artemisia-DB in enabling researchers to uncover new regulatory networks, generate hypotheses, and identify candidate genes for experimental validation. By providing an interactive platform for genomic, transcriptomic, and functional data analysis, the database serves as a valuable resource for advancing our understanding of *A. annua*’s metabolic pathways and regulatory mechanisms.

### Overview of the Artemisia-DB interface

Artemisia-DB provides an integrated, user-friendly platform for exploring the transcriptome landscape of *A. annua*. The interface includes core modules such as Gene Expression, Transcription Factors, Functional Annotation, CRISPR and Epigenetics, JBrowse Genome Browser, BLAST, and Download. Users can visualize expression patterns, explore transcription factor families, access multiple annotation systems, identify gene editing targets, perform sequence alignment, and download processed data.

A comprehensive user guide detailing all features—including tools for co-expression analysis, GO enrichment, and gene sequence retrieval—is provided in the About section of the website and included as supplementary table 2.

## Methods

### Data Sources and Preprocessing

For building the Artemisia-DB, SRA files of RNA-Seq data available on the NCBI’s Sequence Read Archive (SRA), National Genomics Data Center (https://ngdc.cncb.ac.cn/) and Global Pharmacopeia Genome Database (http://www.gpgenome.com/) databases up to 2024-02-05 along with the latest available genome of *A. annua* [39] were downloaded (Supplementary table 1). Gene expression profiling was conducted using a combination of paired-end and single-end short-read RNA-seq data. To improve the accuracy of our transcriptome assembly, we also leveraged long-read sequencing data, which, while typically employed for genome assembly, proved invaluable in resolving complex transcript structures.

Conversion of SRA files to FASTQ was performed using fastq-dump (3.1.0), which is part of the SRA Toolkit. To ensure the quality of the FASTQ files, fastp (0.23.2) (Chen et al., 2018) was used, and FASTQ files with a mean read length lower than 40 and/or Q20 rate lower than 80%, and the adapter sequences were removed.

### Transcriptome Assembly and Quality Assessment

To construct a comprehensive transcriptome for *A. annua*, we integrated both short-read and long-read RNA-Seq data. Short-read sequences were aligned using HISAT2 (v2.2.1), while long-read sequences were aligned with Minimap2 (v2.17). Samples with an overall alignment rate below 50% were excluded from further analysis.

For transcript assembly, StringTie (v2.2.1) [11] was used for short-read data, while both StringTie and IsoQuant were employed for long-read data. The assembled transcripts from short-read sequences were merged using the èstringtie --mergè command to generate a unified transcript set.

### Quality of the assembled transcriptomes

The quality of the resulting transcriptomes was evaluated to identify the most comprehensive and accurate assembly for downstream analysis. Four assemblies were compared: the reference transcriptome, the StringTie-assembled transcriptome derived from both short-read and long-read RNA-Seq data, and the IsoQuant-assembled transcriptome. To assess assembly completeness and quality, BUSCO (v5.7.1) [41] was used, which evaluates transcriptome assemblies based on the presence of conserved single-copy orthologs. This initial evaluation provided a foundational understanding of the quality of each assembly, which was essential for developing a scoring system in the Mikado pipeline. By identifying a robust and representative transcriptome, this approach ensured not only a comprehensive assembly but also a framework for prioritizing transcripts and minimizing redundancy in subsequent analyses.

### Clustering Using MMseqs2

While the StringTie-assembled transcriptome exhibited the highest number of complete BUSCOs and the lowest number of missing BUSCOs, it also displayed a high number of duplications. This indicated a significant level of redundancy within the assembly. To address this issue, MMseqs2 (Many-against-Many sequence searching tool) was employed to cluster highly similar sequences and reduce redundancy [42]. By grouping similar transcripts (--min-seq-id 0.8), the clustering process aimed to refine the assembly, ensuring a more accurate representation of unique genes while maintaining the integrity of biologically relevant sequences.

### Splice-site filtering

To improve the accuracy of splice-site detection in the long-read assemblies, Portcullis was employed. Portcullis is a specialized tool designed to identify and filter splice junctions with high precision, addressing errors commonly found in transcriptome assemblies generated from long-read sequencing data [43]. By applying this filtering step, erroneous or low-confidence splice sites were removed, resulting in a cleaner and more reliable set of splice junctions. This refinement was crucial for ensuring the quality of the transcriptome assembly and for minimizing false positives that could impact downstream analyses, including transcript annotation and functional characterization.

### Merging annotations using the Mikado pipeline

To integrate and refine the four transcriptome assemblies—reference transcriptome, StringTie (short-read and long-read), and IsoQuant—we used the Mikado pipeline [13]. Mikado is designed to identify the most accurate and representative set of transcripts from multiple assembly methods. The pipeline first defines gene loci based on overlap criteria and evaluates each transcript within a locus using up to 50 metrics, including ORF and cDNA size, UTR length, and the relative position of the ORF. Mikado also utilizes BLASTX-based protein similarity and can integrate junction confidence data from Portcullis to improve splice-site accuracy. The best-scoring transcripts are selected as primary gene models, while valid splice variants are retained when compatible with the primary isoform. This approach ensures that the final set of transcripts minimizes redundancy, improves gene model quality, and enhances the representation of expressed loci. Mikado’s ability to integrate data from both short-read and long-read technologies allowed us to merge assemblies generated through different approaches, resulting in a more comprehensive and accurate transcriptome for further downstream analysis.

### Gene-level transcript abundance

After generating the representative assembly for *A. annua*, transcript abundances were estimated using the ’mapping-based’ mode of Salmon (v1.10.0) with the ‘--dumpEq’ option, which produces the equivalence classes that can be used in differential transcript usage analysis [44]. Variations in library size across samples are taken into consideration by transcripts per million (TPM). However, TPM does not take variations in transcript length into consideration, meaning that, even when transcripts are expressed at the same level, longer transcripts will typically have higher TPM values than shorter transcripts. To obtain the gene-level transcript abundances, the bias-corrected counts without an offset method from tximport R package was used [45]. The "lengthScaledTPM" argument in tximport adjusts for differences in both library size and transcript length across samples. For the transcript-level transcript abundance estimates, the “dtuScaledTPM” argument was used, which first scales using the median transcript length among isoforms of a gene, and then by library size.

### Dimensionality reduction

Because bioinformatics data contains numerous attributes (variables), they are naturally high-dimensional. This high dimensionality can present major hurdles for data analysis, obscuring valuable biological insights and complicating downstream analysis. Therefore, appropriate procedures should be taken to boost the biological significance of the data by reducing the number of attributes using dimensionality reduction techniques [46]. For transcriptomics in particular, dimensionality reduction helps to mitigate noise and enhance the signal from the biological data. To achieve this goal, techniques such as principal component analysis (PCA), t-distributed stochastic neighbor embedding (t-SNE) [47], and uniform manifold approximation and projection (UMAP) [48] are commonly employed. In the present study, we used scran R package [49]. After log-transforming the gene counts normalized by library size, we extracted the top 3000 genes with the highest biological components to model the mean-variance relationships. Following performing PCA analysis, 11 principal components, which accounted for 70% of the variation, were selected for dimensionality reduction using the t-SNE and UMAP algorithms. Six different perplexity values (5, 10, 15, 20, 25, 30) for t-SNE and six different numbers of nearest neighbors (10, 20, 30, 40, 50, 60) for UMAP were tested. The optimal values were determined based on visual inspection.

### Identification of broadly expressed and body part-specific genes

Identifying broadly expressed and tissue-specific genes is crucial for understanding gene function and regulation. Broadly expressed genes often play essential roles in basic cellular processes, while body part-specific genes are typically involved in specialized tissue functions. Analyzing these expression patterns helps researchers understand gene functions and regulatory mechanisms [50]. To achieve this, we calculated the tissue specificity index (τ index) for each gene by using a log-transformed matrix of the median TPM values per body part (Yanai et al., 2005). Based on the criteria established by Lüleci and Yılmaz (2022), genes were categorized into four groups according to their τ index and median TPM values: (1) Null expression: genes with median TPM < 1 in all tissues, indicating no expression; (2) Weak expression: genes with median TPM < 5 in all tissues; (3) Broad expression: genes with τ < 0.85 and median TPM > 5; and (4) Body part-specific expression: genes with τ ≥ 0.85 and median TPM > 5.

### Functional annotation of the *A. annua* transcriptome

To create a unified database for the functional annotation of the *A. annua* transcriptome, several analyses were performed. Transcription factors (TFs) were identified using two approaches: first, by performing a BLASTx search against a collection of TFs from the PlantTFDB [51], and second, by extracting PFAM IDs related to 63 different TF families through InterProScan. For the BLASTx analysis, Diamond was used with the following parameters: --ultra-sensitive --id 60 --evalue 1e-5 --min-score 150 [52]. EggNOG-mapper (2.1.12) [53] using the HMMER3 mode was used to annotate orthologous groups and functional categories. InterPro domains, Gene Ontology (GO) terms, and KEGG pathways were assigned using InterProScan (5.67- 99.0); The KEGG ORTHOLOGY (KO) database (https://www.genome.jp/kegg/ko.html) was used to obtain KO descriptions. Additionally, BLASTx against *Arabidopsis thaliana* protein-coding genes (Araport11 release, 2022-09-15) and the Uniprot database (release 01.03.2023) was performed to further enhance the functional characterization of the transcriptome. CRISPR guide sequences were identified using CRISPRCasFinder (version 4.2.30) [54]. A list of artemisinin-related genes was obtained from recently published research [2]. Their sequences were retrieved from the NCBI database, and the corresponding gene IDs in the current annotation were identified using BLASTn for further analysis. For enrichment analysis, enricher() function of the clusterProfiler package [55] was utilized, with all expressed genes serving as the background set.

### Development of the web application

The web application was developed using the Shiny framework [56], with a Bootstrap-based layout. A MySQL database was used to manage data associated with various sections of the application, ensuring well-organized data retrieval and storage. A partitioned Parquet directory was employed to store the expression database, with the arrow R package [57] providing an efficient interface between R and the Apache Arrow platform. For data visualization, the application leveraged the ggplot2 [58], plotly [59], and ComplexHeatmap [60] packages.

## Author Contribution

**A.T.** conceived the study, performed all data analyses, developed the web application, and wrote the manuscript. **F.A.-S.** contributed to the data analysis, manuscript revision, and discussion. **X.F.** and **K.T.** provided funding support. **K.T.** supervised the study and is the corresponding author. **Y.Z., L.L., and Y.W.** contributed to manuscript revision and discussion. All authors approved the final manuscript.

## Acknowledgments

We acknowledge the contributions of the researchers and organizations who generated the genome and RNA-sequencing data utilized in this study. This work was supported by the National Natural Science Foundation of China (82274047) and the Bill & Melinda Gates Foundation (INV-027291). Under the grant conditions of the Bill & Melinda Gates Foundation, a Creative Commons Attribution 4.0 Generic License has already been assigned to the Author Accepted Manuscript version that might arise from this submission. The computations in this paper were run on the Siyuan-1 cluster supported by the Center for High Performance Computing at Shanghai Jiao Tong University.

## Data Availability Statement

The datasets presented in this study are available in the mentioned online repositories. The code used for analysis is available at the GitHub repository: https://github.com/ayattaheri/artemisia-db.

---

**Figure S 1.**
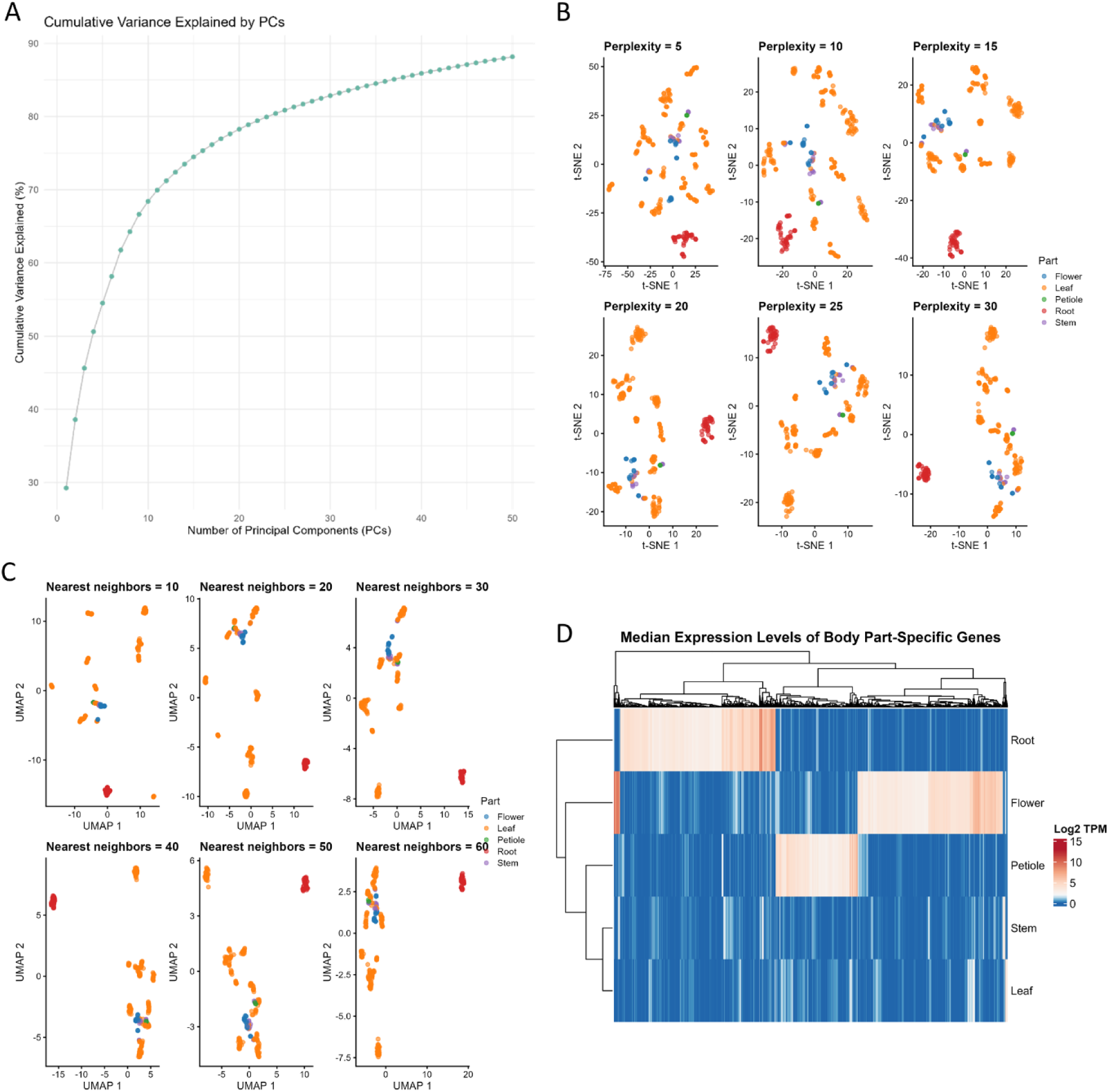
(A) Cumulative variance explained by each principal component (PC) in the transcriptome analysis. (B) t-SNE plots generated using different perplexity values (5, 10, 15, 20, 25, 30). (C) UMAP plots based on varying numbers of nearest neighbors (10, 20, 30, 40, 50, 60). (D) Heatmap of median expression levels of tissue-specific genes across different tissues.

**Figure S 2.**
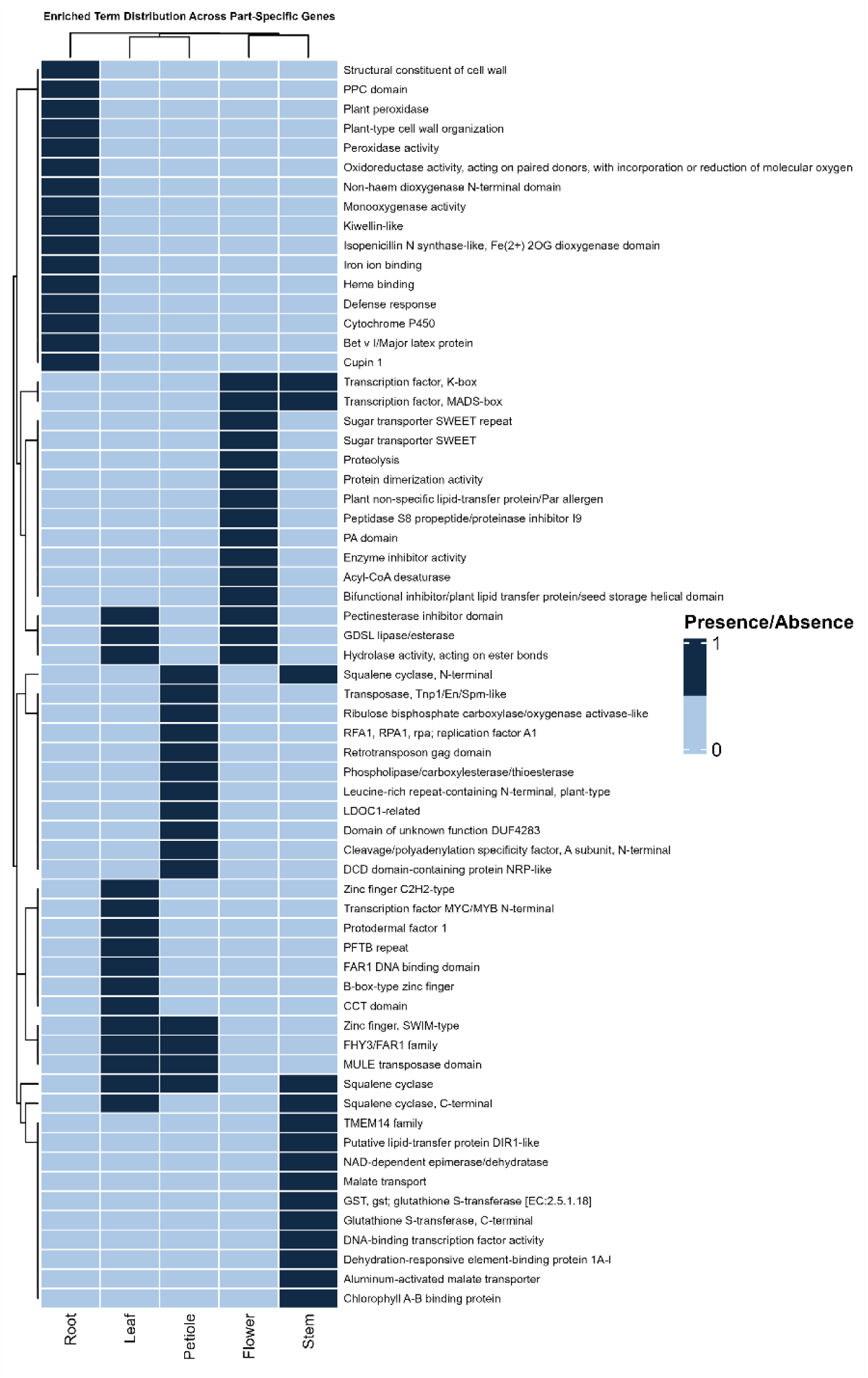
Heatmap of Gene Ontology (GO) terms and InterPro domain enrichment for tissue-specific genes.

**Figure S 3.**
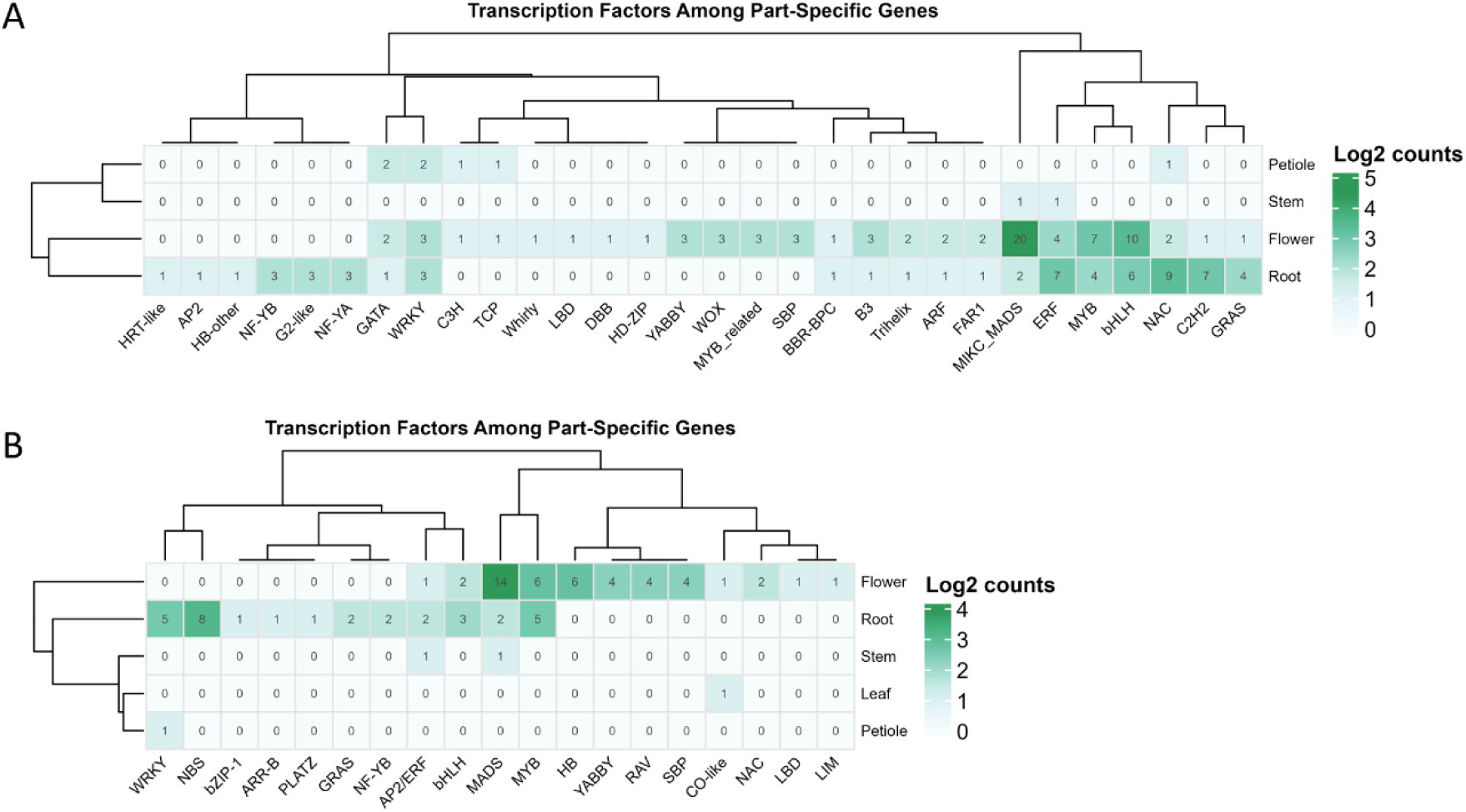
Heatmap of transcription factor distribution across tissue-specific genes identified from PlantTFDB. (B) Heatmap of transcription factor distribution across tissue-specific genes identified using Pfam IDs.

