## Supplementary Table 2 for "Artemisia Database: A Comprehensive Resource for Gene Expression and Functional Insights in *Artemisia annua*"

### Tutorial: Exploring the Artemisia Database

Welcome to the **Artemisia Database**, a comprehensive platform designed to facilitate research on *Artemisia annua* gene expression, transcription factors, functional annotations, and more. This tutorial will walk you through the database's key features, showing you how to navigate its interface and utilize its tools effectively. Whether you're a biologist, bioinformatician, or researcher, this guide will help you get started.

---

#### Table of Contents


1. Getting Started
2. Home Page Overview
3. Gene Expression
   - Global Expression Viewer
   - Median Expression Per Part
   - Artemisinin Pathway Genes
   - Overall Category Distribution
4. Transcription Factors
   - PlantTFDB
   - Pfam
   - Tissue-specific TFs
5. Functional Annotation
6. Gene Editing and Epigenetics
   - CRISPR
   - Methylation
7. Tools
   - JBrowse
   - BLAST
   - Co-expression Analysis
   - GO Enrichment
   - Get Gene Sequences
8. Download
9. About the Database
10. Tips and Troubleshooting


---

#### Getting Started

##### Accessing the Database

- Open your web browser and navigate to the Artemisia Database URL (`https://artemisia-db.com`).
- The interface is built using Shiny, so it's interactive and user-friendly.

---

#### Home Page Overview

Upon loading the database, you'll land on the **Home** tab:

- **Welcome Message**: A brief introduction to the database and its purpose—exploring *Artemisia annua* gene expression data.
- **Flowchart**: A visual representation of how the database was developed, centered on the page for easy reference.

---

### Gene Expression

The **Gene Expression** menu contains four powerful tools for visualizing expression data.

#### Global Expression Viewer

- **Purpose**: View a t-SNE plot of gene expression colored by plant body parts and explore associated metadata.

- **Steps**:

    1. Navigate to **Gene Expression > Global Expression Viewer**.
    2. Observe the interactive t-SNE plot (powered by Plotly) showing expression clusters.
    3. Scroll down to the "Filtered t-SNE Metadata" table to see detailed sample information.

- **Example**: Hover over points in the t-SNE plot to identify clusters representing samples from specific plant parts, such as "Leaf" or "Root." Click on any point to filter the "Filtered t-SNE Metadata" table below based on your selection. This table provides detailed information about the chosen sample, including its metadata. If a sample is linked to a published study, the `Pubmed_ID` column will contain a hyperlink—click it to visit the corresponding PubMed page for more details about the research.

t-SNE Colored by Body

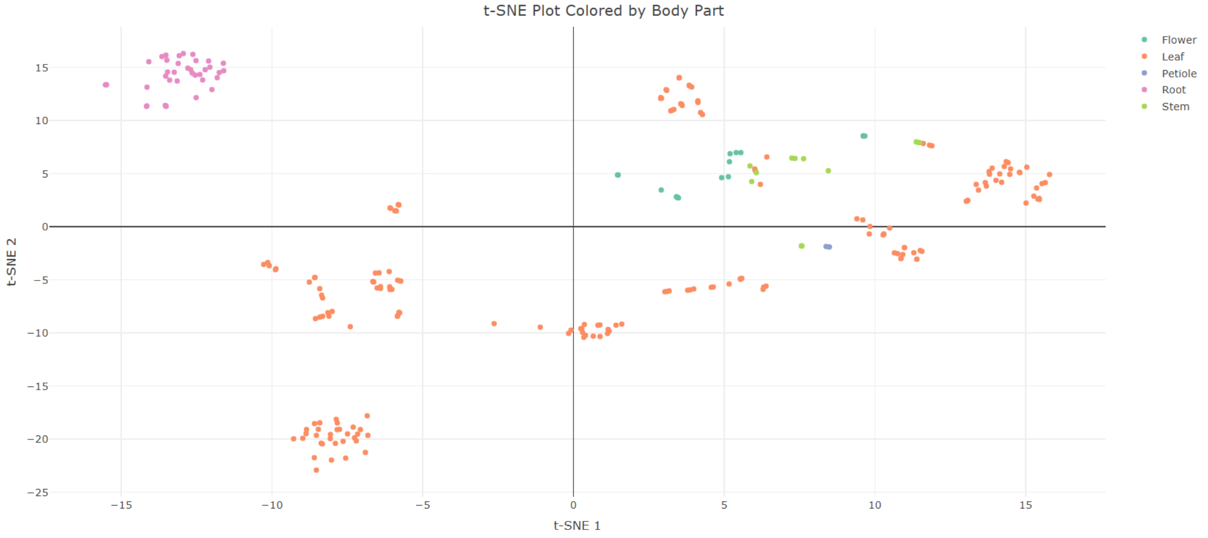


Filtered t-SNE Metadata

Show 10 entries

Search:

| | Run | Project | Ecotype | Part | AGE | Pubmed_ID |
|---|---|---|---|---|---|---|
| 1 | ARTAN0001 | PRJCA009719 | LQ-9 | Root | 4-month-old plant | 35655434 |
| 2 | ARTAN0002 | PRJCA009719 | LQ-9 | Root | 4-month-old plant | 35655434 |
| 3 | ARTAN0003 | PRJCA009719 | LQ-9 | Root | 4-month-old plant | 35655434 |
| 4 | ARTAN0004 | PRJCA009719 | HAN1 | Root | 4-month-old plant | 35655434 |
| 5 | ARTAN0005 | PRJCA009719 | HAN1 | Root | 4-month-old plant | 35655434 |

*Figure 1: The t-SNE plot displays gene expression data, with points colored by plant parts like Leaf and Root. Hover over or click points to interact with the data.*

#### t-SNE Colored by Body

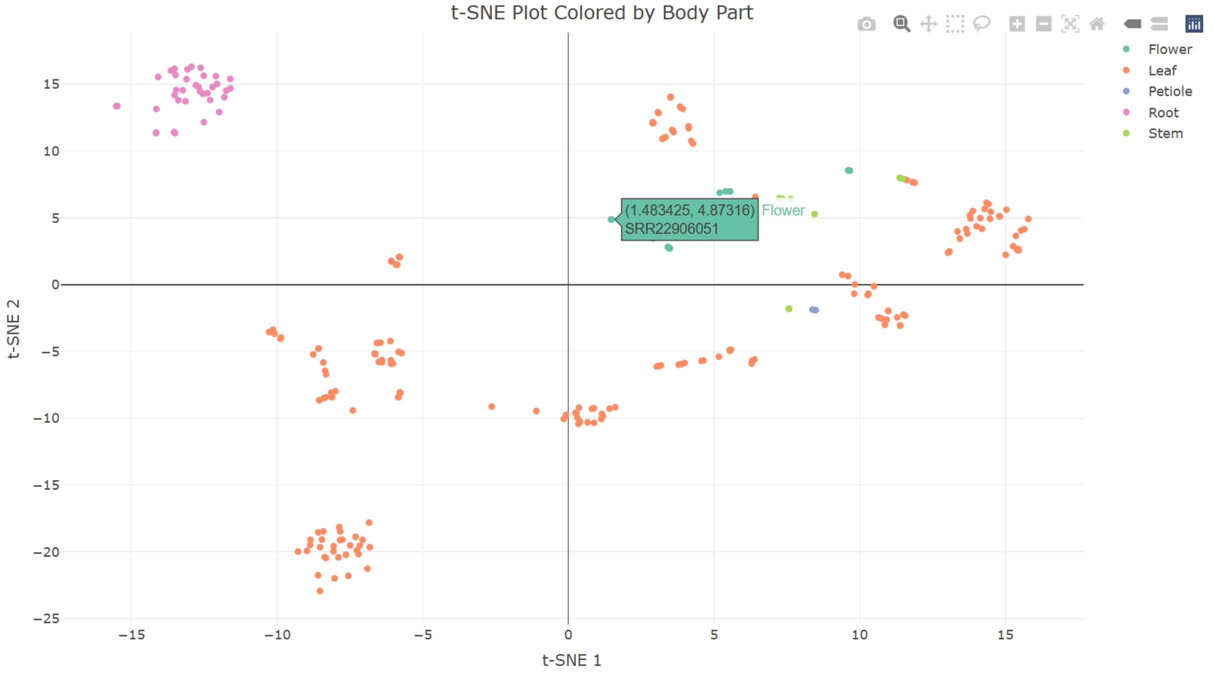


#### Filtered t-SNE Metadata

Show 10 entries Search:

| | Run | Project | Ecotype | Part | AGE | Pubmed_Link |
|---|---|---|---|---|---|---|
| 254 | SRR22906051 | PRJNA916469 | Huhao 1 | Flower | 6-week-old | 37239820 |

*Figure 2: The Filtered t-SNE Metadata table updates based on your selection, showing sample details and a clickable Pubmed_ID link.*

---

##### Median Expression Per Part

- **Purpose**: Calculate and visualize median gene expression across selected plant parts.
- **Steps**:
  1. Go to **Gene Expression > Median Expression Per Part**.
  2. Enter up to 100 gene IDs (one per line) in the text box or upload a `.txt`/`.csv` file. We added the example gene ID buttons to auto-fill the text box.
  3. Select plant parts (e.g., "Root", "Leaf") using the checkboxes.
  4. Click **Calculate Median Expression**.
  5. View the resulting heatmap (log2-transformed data).
  6. Download the plot in PNG format via clicking on the camera icon on the top right corner of the plot.
- **Example**: Enter `mikado.chr7G1274` and `mikado.chr4G1337`, select "Leaf" and "Flower", then click "Calculate" to see a heatmap comparing their expression.

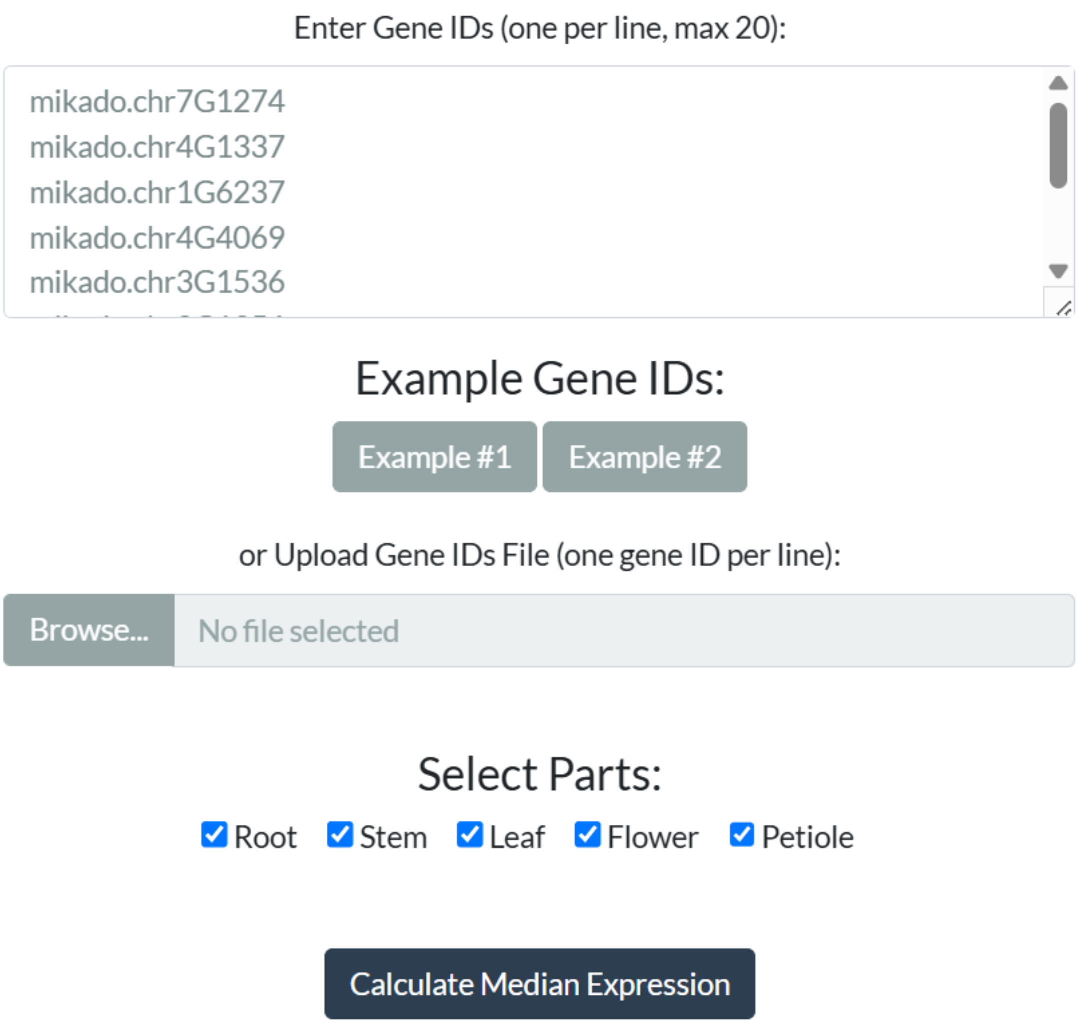


*Figure 3: Clicking an example gene ID button (e.g., "Example#1") auto-fills the text area with gene IDs. Select different plant parts (e.g., Leaf, Flower) to analyze their median expression in the heatmap.*

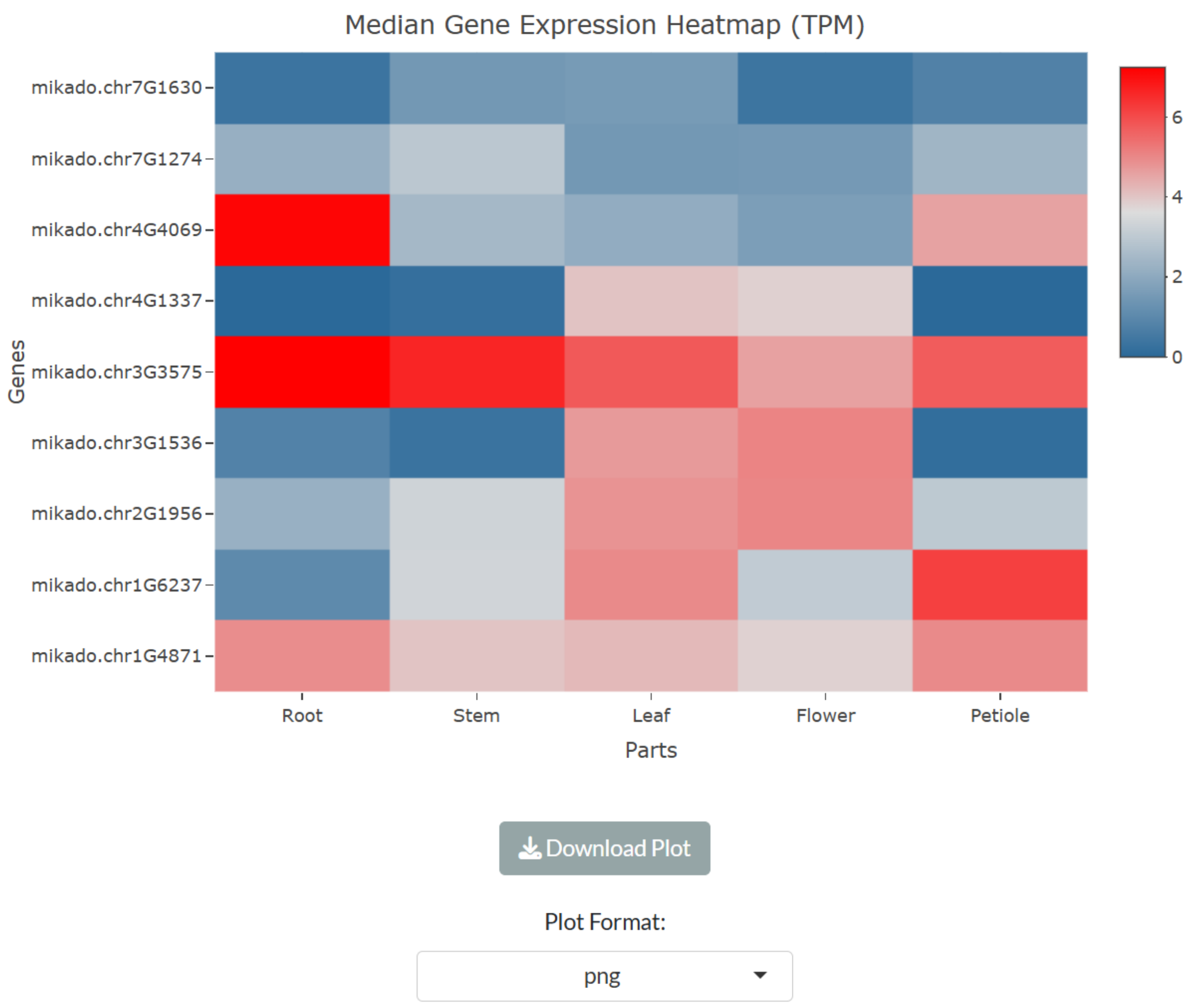


*Figure 4: The heatmap displays the median expression (log2-transformed) of example gene IDs across selected plant parts, generated after clicking 'Calculate Median Expression'.*

---

#### Artemisinin Pathway Genes

- **Purpose**: Focus on genes linked to artemisinin production in *Artemisia annua*, providing detailed information through a table with tabs for main data, expression, and sequences.
- **Steps**:
    1. Navigate to **Gene Expression > Artemisinin Pathway Genes**.
    2. View the "Main Table" tab, which lists artemisinin-related genes and their details.
    3. Search by gene names or gene IDs (one per line) in the search box to filter the table and view specific information:
    4. For gene IDs, enter the ID and press the **Search by Gene ID** button.
    5. For gene names, enter the name and press the **Search by Gene Name** button.
    6. Select one or more rows in the "Main Table" to reveal the **Check Gene Expression** and **Copy Gene ID(s)** buttons at the bottom left.
    7. Click **Copy Gene ID(s)** to copy the selected gene IDs to the clipboard for use in other analyses.
    8. Click **Check Gene Expression** to switch to the "Expression" tab, where you can select plant parts (e.g., "Leaf", "Root") and click **Calculate Median Expression** to generate a heatmap.

9. Switch to the "Sequence" tab to view and download the sequences of the selected genes as a text file.

---

##### Overall Category Distribution

- **Purpose**: Provide an overall view of gene categories in *Artemisia annua* (e.g., specific, broad) using a donut chart, allowing users to explore classifications and related data.
- **Steps**:
    1. Navigate to **Gene Expression > Overall Category Distribution**.
    2. View the donut chart displaying gene categories (e.g., "Specific").
    3. Click a segment (e.g., "Specific") to filter the table below by that category.
    4. Alternatively, enter gene ID(s) in the search box to find its category and display its details in the table.
    5. Select one or more rows in the "Main Table" to reveal the **Check Gene Expression** and **Copy Gene ID(s)** buttons at the bottom left.
    6. Click **Copy Gene ID(s)** to copy the selected gene IDs to the clipboard for use in other analyses.
    7. Click **Check Gene Expression** to switch to the "Expression" tab, where you can select plant parts (e.g., "Leaf", "Root") and click **Calculate Median Expression** to generate a heatmap.
    8. Switch to the "Sequence" tab to view and download the sequences of the selected genes as a text file.

---

#### Transcription Factors

The **Transcription Factors** menu offers three sub-tabs for analyzing transcription factors (TFs).

##### PlantTFDB

- **Purpose**: Explore transcription factor (TF) families from the PlantTFDB database.

- **Steps**:

    1. Navigate to **Transcription Factors > PlantTFDB**.
    2. View the interactive TF family distribution plot.
    3. Click on any bar in the plot (e.g., "bZIP") to filter the "Main Table" below based on your selection. You can also search different TF families based on the gene IDs.
    4. Switch to the "Details" box and explore:
        - **Main Table**: A filterable table displaying TF data. Select one or more rows to reveal the **Check Gene Expression** and **Copy Gene ID(s)** buttons at the bottom left. Clicking **Check Gene Expression** button automatically switches you to the "Expression" tab with the selected gene IDs pre-filled. Click **Copy Gene ID(s)** to copy the selected gene IDs to the clipboard for use in other analyses.
        - **Expression**: On this tab, choose different plant parts (e.g., "Root", "Leaf"), then click **Calculate Median Expression** to generate a heatmap of gene expression for the selected genes.
        - **Sequence**: On this tab, view and download the transcript sequences for your selected genes as a text file.

- **Example**: Start by clicking the "bZIP" bar in the TF family distribution plot to filter the table for bZIP family TFs. In the "Main Table," select rows for specific genes (e.g., two bZIP TFs), then click **Check Gene Expression**. On the "Expression" tab, select "Leaf" and "Stem," click **Calculate Median Expression**, and view the resulting heatmap, which you can download in your preferred format.

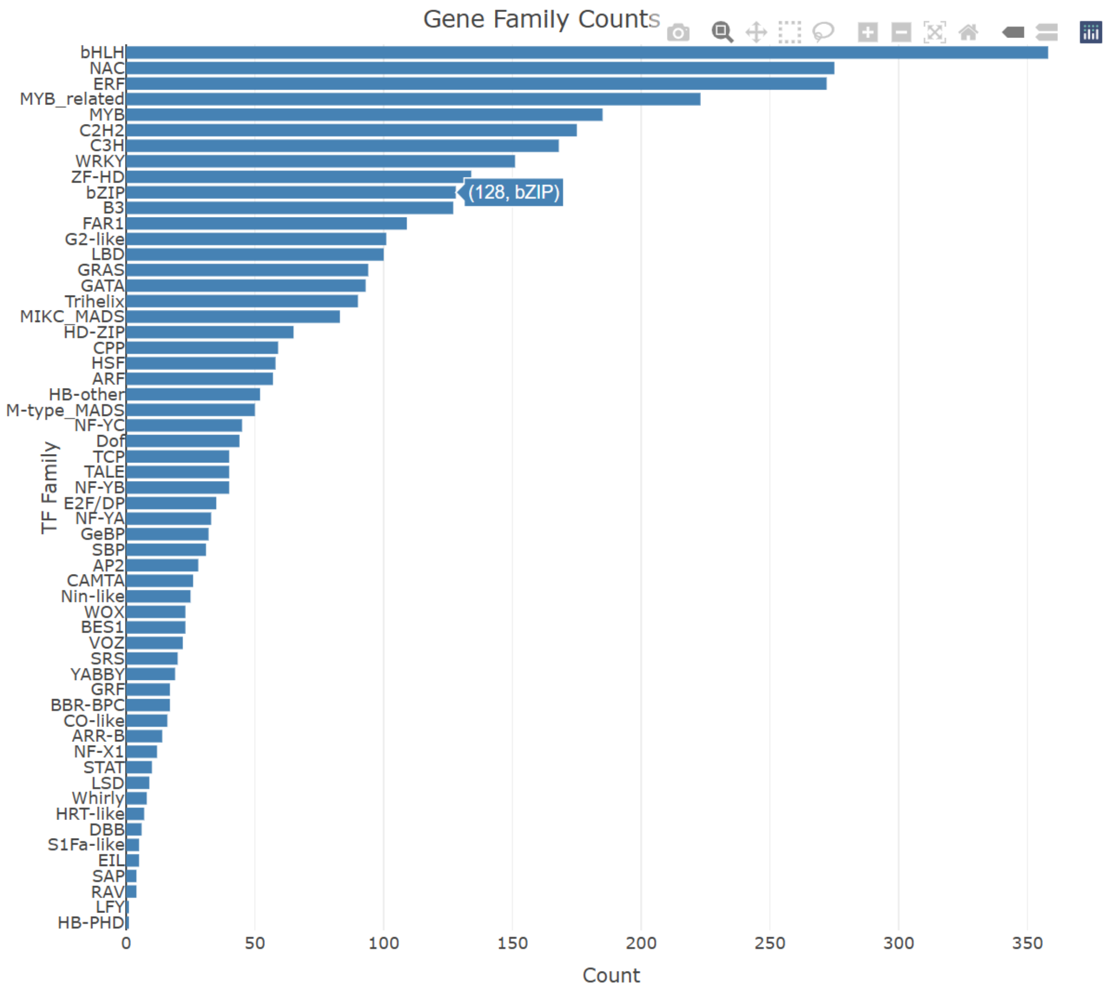


*Figure 5: The TF family distribution bar plot shows the number of TFs per family. Clicking the "bZIP" bar filters the table below to display only bZIP family transcription factors.*

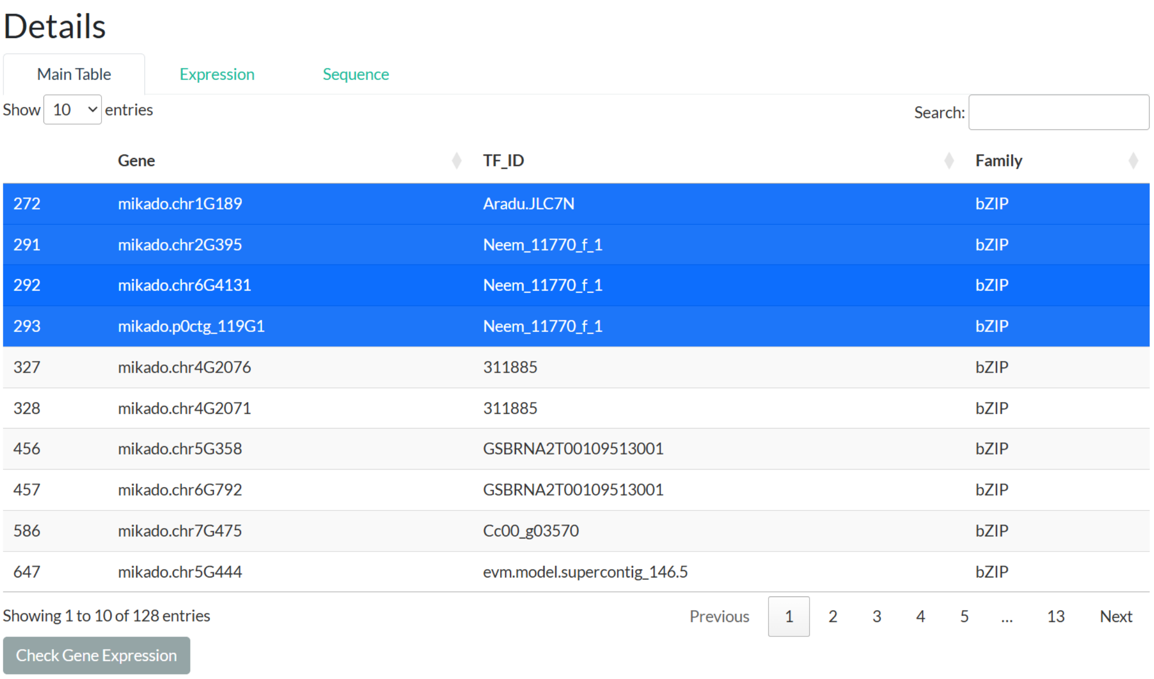


*Figure 6: The Main Table is filtered to show bZIP family TFs after clicking the "bZIP" bar. Highlighted rows indicate selected genes, and the "Check Gene Expression" button appears at the bottom left, ready to switch to the Expression tab.*

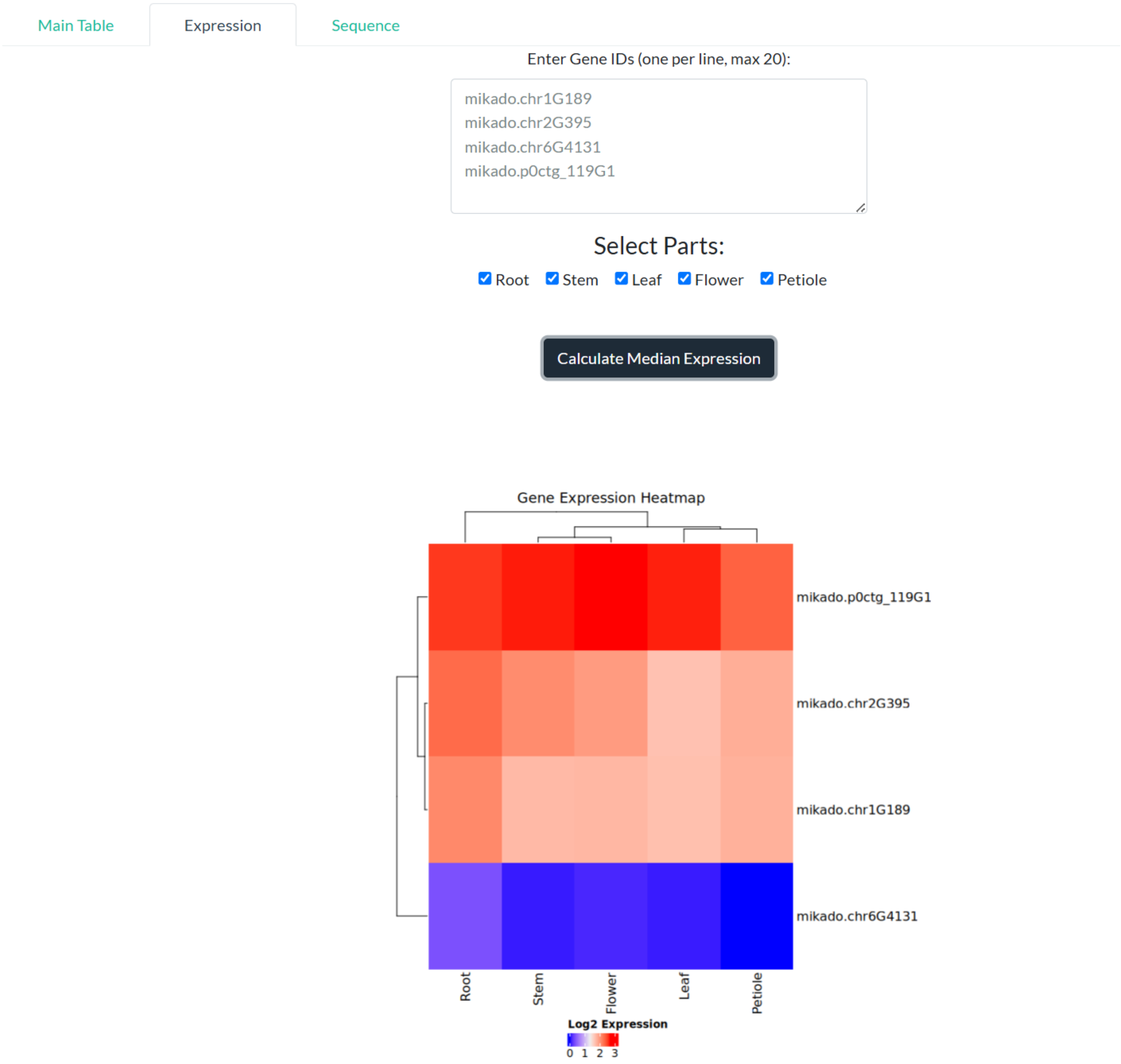


*Figure 7: On the Expression tab, select plant parts (e.g., Leaf, Stem) for the chosen bZIP genes, then click "Calculate Median Expression" to display the heatmap. Use the "Download Plot" button to save it as PNG, JPEG, or PDF.*

##### Pfam

- **Purpose**: Analyze transcription factors (TFs) based on Pfam domain annotations.
- **Steps**: Similar to PlantTFDB, with a focus on Pfam domains instead of PlantTFDB families. Navigate to **Explore Transcription Factors > Pfam**, view the family distribution plot, filter the table by clicking bars, and explore the "Details" box (Main Table, Expressions, Sequence).

##### Tissue-specific TFs

- **Purpose**: Investigate TFs specific to certain plant tissues, such as Leaf or Root.

- **Steps**:

    1. Navigate to **Transcription Factors > Tissue-specific TFs**.
    2. View the heatmap plot showing TF expression across tissues.
    3. Click a heatmap cell (e.g., CO-like TF in Leaf) to filter the "Main Table" in the "Details" box below.

4. Explore the "Details" box:
    - **Main Table**: Filterable table of tissue-specific TF data. Select rows to enable the **Check Gene Expression** button.
    - **Expression**: Calculate and view a heatmap for selected genes across plant parts.
    - **Sequence**: Access and download sequences for selected TFs.

- **Example**: In the "Tissue-specific TFs" heatmap, click the cell for the CO-like TF specific to Leaf tissue to filter the table below. Select the CO-like TF row in the "Main Table," click **Check Gene Expression**, and on the "Expression" tab, generate a heatmap for "Leaf" and "Root" to compare expression patterns.

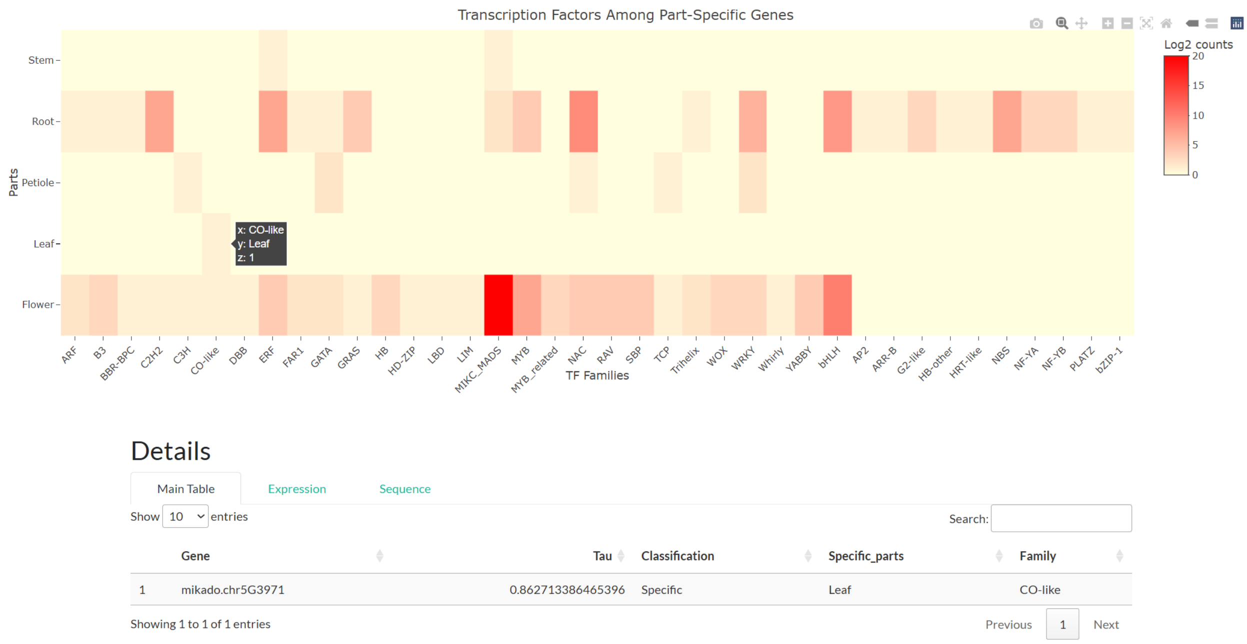


*Figure 8: The heatmap shows tissue-specific TFs, with the CO-like TF cell for Leaf tissue clicked, filtering the table below to display only Leaf-specific CO-like TFs.*

---

### Functional Annotation

- **Purpose**: Query functional annotations of *Artemisia annua* genes using three distinct methods.

- **Steps**:

1. Navigate to **Functional Annotation**.
2. Choose one of three query options:
    - **Annotation Table**: Select a database (e.g., "GO") from the dropdown and enter annotation IDs in the text area. By default, example IDs are provided for each database (e.g., GO, Pfam, KEGG) to guide users—enter similar IDs matching the selected database's format.
    - **Gene-Based Queries**: Input any gene IDs of interest based on the latest gene model developed for this database. Results will appear in a table with multiple tabs, each representing a different annotation database (e.g., GO, Pfam, KEGG).
    - **Functional Descriptions**: Search using keywords (e.g., "RNA polymerase") in the text input field. Results are displayed in a table format similar to the gene-based query, with tabs for different annotation types.
3. Click the corresponding **Search** button for your chosen query method.

4. View the results in dynamically generated tabs below the query section.

- **Example**: For a gene-based query, enter a gene ID (e.g., `mikado.Super-Scaffold_100038G100`) in the "Gene-Based Queries" text area and click **Search Gene IDs**. The results will display in a table with multiple tabs, such as GO, Pfam, and KEGG, showing annotations linked to `mikado.Super-Scaffold_100038G100`.

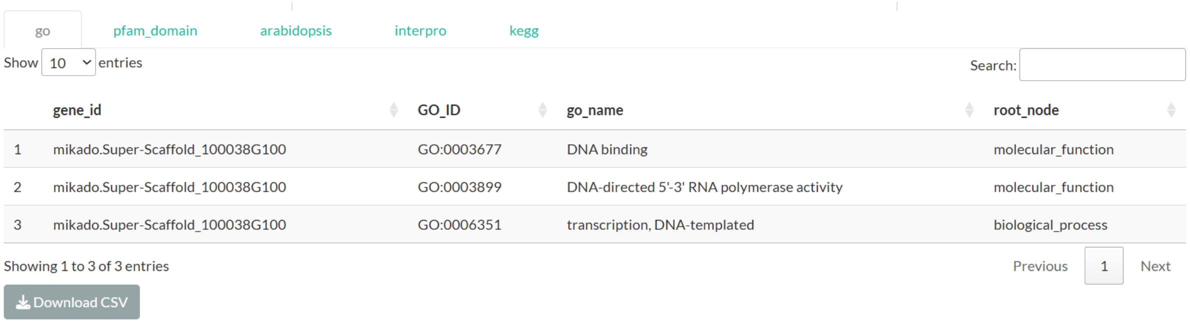

go pfam_domain arabidopsis interpro kegg

Show 10 entries Search:

| | gene_id | GO_ID | go_name | root_node |
|---|---|---|---|---|
| 1 | mikado.Super-Scaffold_100038G100 | GO:0003677 | DNA binding | molecular_function |
| 2 | mikado.Super-Scaffold_100038G100 | GO:0003899 | DNA-directed 5'-3' RNA polymerase activity | molecular_function |
| 3 | mikado.Super-Scaffold_100038G100 | GO:0006351 | transcription, DNA-templated | biological_process |

Showing 1 to 3 of 3 entries Previous 1 Next

Download CSV

*Figure 9: After searching for a gene ID (e.g., GeneID1), the results appear in a table with multiple tabs (e.g., GO, Pfam, KEGG), each displaying different annotations for the gene.*

---

### Gene Editing and Epigenetics

#### CRISPR

- **Purpose**: Explore gene annotations and CRISPR-related sequences for *Artemisia annua*, enabling users to investigate gene editing potential.
- **Steps**:
  1. Navigate to **Gene Editing and Epigenetics > CRISPR**.
  2. View the table of CRISPR-related gene annotations.
  3. Enter one or more gene IDs in the search box to filter the table and check for associated CRISPR sequences.
  4. Select one or more rows in the filtered table to reveal the **Check Gene Expression** and **Copy Gene ID(s)** buttons at the bottom left.
  5. Click **Copy Gene ID(s)** to copy the selected gene IDs to the clipboard for use in other analyses.
  6. Click **Check Gene Expression** to switch to the "Expression" tab with the selected gene IDs pre-filled.
  7. On the "Expression" tab, choose plant parts (e.g., "Leaf", "Root"), then click **Calculate Median Expression** to generate a heatmap.
  8. Switch to the "Sequence" tab to view and download the sequences of the selected genes as a text file.

#### Methylation

- **Purpose**: Analyze methylation data (e.g., N6-methyladenosine (m6A), 5-methyladenosine) for *Artemisia annua* genes to explore epigenetic modifications.
- **Steps**:
  1. Navigate to **Gene Editing and Epigenetics > Methylation**.
  2. Select a methylation type from the dropdown (e.g., "N6-methyladenosine (m6A)").
  3. Choose a subfilter (e.g., "Writer", "Reader", "Eraser") to refine the dataset.

4. Enter one or more gene IDs in the search box to filter the table and view specific methylation-related annotations.
5. View the table of methylation-related gene annotations.
6. Select one or more rows in the filtered table to reveal the **Check Gene Expression** and **Copy Gene ID(s)** buttons at the bottom left.
7. Click **Copy Gene ID(s)** to copy the selected gene IDs to the clipboard for use in other analyses.
8. Click **Check Gene Expression** to switch to the "Expression" tab with the selected gene IDs pre-filled.
9. On the "Expression" tab, choose plant parts (e.g., "Leaf", "Root"), then click **Calculate Median Expression** to generate a heatmap.
10. Switch to the "Sequence" tab to view and download the sequences of the selected genes as a text file.

---

### Tools

#### JBrowse

- **Purpose**: Visualize genomic data for *Artemisia annua* using JBrowse, based on the latest annotation and sequence files developed in this study.

- **Steps**:

  1. Navigate to **Tools > JBrowse**.
  2. Click the **Open** button to access the genome browser.
  3. Click **Open Track Selector** in the sidebar to view available tracks.
  4. Select the "Artemisia Annotation" (GFF3) and "Genome Sequence" (FASTA) files to load them into the browser.
  5. Zoom in on a region of the genome to view gene features, such as exons, introns, or UTRs.
  6. Click on any feature (e.g., a gene or CDS) to open a new window with detailed information.
  7. In the feature window:
     - Click **Show Feature Sequence** to display and copy the sequence of the selected feature.
     - Use the dropdown menu to extract the sequence with default 1000 bp upstream and downstream.
     - Click the wheel icon next to the dropdown to modify the extraction range (e.g., change to 500 bp or 2000 bp).

- **Example**: After opening JBrowse and loading the Artemisia annotation and genome sequence tracks, zoom into a region to see gene features from the GFF3 file. Select a gene feature to open its details window, then click **Show Feature Sequence** to view its sequence. From the dropdown menu, choose to extract 1000 bp upstream and downstream, and adjust this to 1500 bp by clicking the wheel icon and modifying the range.

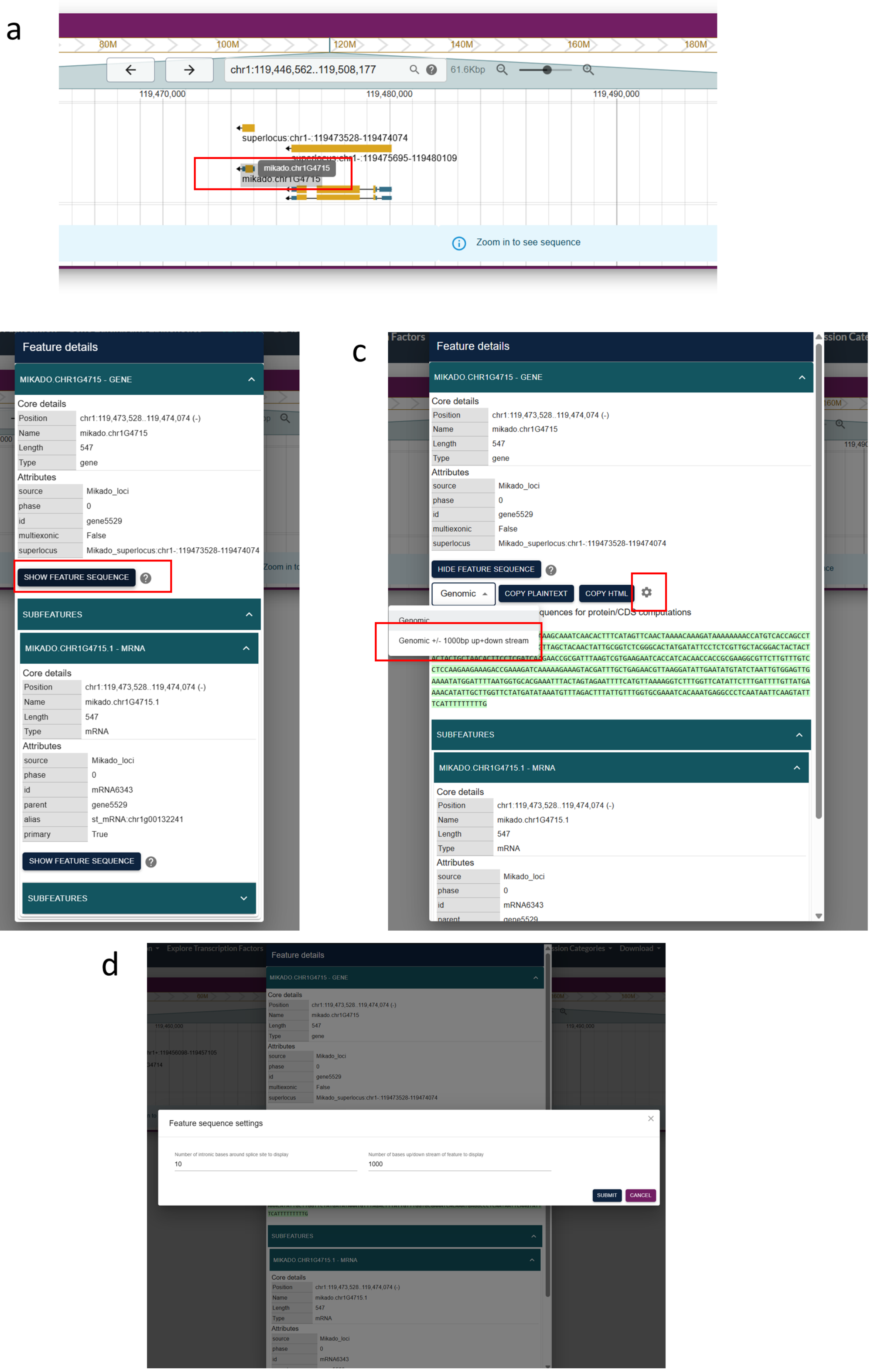


*Figure 10: Exploring features in JBrowse.*
*(a) The JBrowse interface displays various gene features after loading the Artemisia annotation and genome sequence tracks and zooming in. (b) Clicking a feature opens a new window with detailed information. (c) Selecting "Show Feature Sequence" reveals the sequence, with a dropdown menu to extract 1000 bp upstream and downstream. (d) Clicking the wheel icon allows modification of the extraction range (e.g., to 1500 bp).*

### BLAST

- **Purpose**: Perform sequence similarity searches to identify corresponding *Artemisia annua* gene IDs based on the latest genome annotation. This tool is ideal for users who have a gene sequence from another species or a gene ID from previous *Artemisia* annotation files and want to find its match in the database, unlocking access to all features (e.g., expression, annotations, sequences).

- **Steps**:

  1. Navigate to **Tools > BLAST**.
  2. Enter a FASTA-formatted sequence (e.g., from another species) or upload a file (<2MB) containing the query sequence.
  3. Adjust settings as needed:
     - **Minimum Identity**: Set the similarity threshold (default: 80%).
     - **E-value**: Set the statistical significance threshold (default: 1e-5).
  4. Click **Run BLAST** to search the sequence against the *Artemisia annua* genome and latest gene IDs.
  5. View the results table, which includes matching *Artemisia* gene IDs, and download it as a CSV file for further use.

- **Example**: Suppose you have a gene sequence from another species, such as `>Seq1\nATCGATCG...`, or a gene ID from an older *Artemisia* annotation. Input this sequence in the text box (e.g., `>Seq1\nATCGATCG...`), adjust the E-value to 1e-5, and click **Run BLAST**. The results will list the corresponding *Artemisia annua* gene ID (e.g., `GeneID1`) from the latest annotation, which you can then use to explore expression, functional annotations, or sequences across the database.

---

#### Co-expression Analysis

- **Purpose**: Identify and analyze co-expressed genes in *Artemisia annua* across selected bioprojects and plant parts, with options for correlation methods and enrichment analysis.
- **Steps**:
  1. Navigate to **Tools > Co-expression Analysis**.
  2. Enter one or more gene IDs in the top-left search box to query co-expressed genes.
  3. Select a correlation method ("Pearson" or "Spearman") from the dropdown menu.
  4. Choose one or more bioprojects from the "Filter by Bioproject" list, or select "ALL" to include all projects.
  5. Select plant parts (e.g., "Leaf", "Root") from the "Select Plant Part(s)" list, which updates based on the chosen bioprojects.
  6. Adjust the "FDR Rate Threshold" and "Minimum Absolute Correlation" (0 to 1) using the sliding menus to filter results.
  7. Click **Run Co-expression** to generate results, displayed in a three-tabbed table:
     - **Table**: Lists co-expressed genes with gene ID, correlation, value, and FDR.
     - **Network Graph**: Visualizes the top 10 co-expressed genes based on user inputs.
     - **GO Enrichment**: Displays a dot plot of enriched terms for co-expressed genes. Hover over the plot to reveal a camera icon for saving as a PNG. Below, a table provides details on enriched terms, filterable by "Root Node" and a "Max P.adjust" slider. Use the **Download** button to save the filtered table.

8. View the metadata table on the right, which updates based on selected bioprojects and plant parts.

---

##### GO Enrichment

- **Purpose**: Perform Gene Ontology (GO) enrichment analysis for a user-defined list of *Artemisia annua* gene IDs to identify enriched biological terms.
- **Steps**:
    1. Navigate to **Tools > GO Enrichment**.
    2. Input a list of gene IDs (one per line) in the text area or upload a text file with one gene ID per line.
    3. Click **Run GO Enrichment** to generate results, displayed in two components:
        - **Dot Plot**: Visualizes enriched GO terms. Hover over the plot to reveal a camera icon for saving as a PNG.
        - **Table**: Lists enriched terms with details, filterable by "Root Node" and a "Max P.adjust" slider. Use the **Download** button to save the filtered table.

---

##### Get Gene Sequences

- **Purpose**: Retrieve and download gene sequences for specified *Artemisia annua* gene IDs.
- **Steps**:
    1. Navigate to **Tools > Get Gene Sequences**.
    2. Enter gene IDs (one per line) in the text area (e.g., `mikado.chr1G1813`).
    3. If ≤5 gene IDs are entered, view the sequences in the output area below the text input.
    4. Click **Download Sequences** to save the sequences as a text file. Note: For >5 gene IDs, sequences are not displayed on-screen but can still be downloaded.

#### Download

- **Purpose**: Access raw data files from the *Artemisia annua* database, including gene expression matrices, project-specific data, and the latest annotation and transcriptome assembly files created in this study.
- **Options**:

    1. **Download by Body Part**

        - **Purpose**: Download gene expression matrices generated in this research, organized by plant body parts.
        - **Steps**:
            1. Navigate to **Download > Download by Body Part**.
            2. Select at least one body part (e.g., "Leaf", "Root") from the available options.
            3. Choose the expression data type:
                - **TPM**: Transcripts Per Million.
                - **Bias-corrected Counts**: Normalized count data.
            4. Select the expression level:
                - **Gene Level**: Expression aggregated at the gene level.
                - **Transcript Level**: Expression at the transcript level.
            5. Click **Search** to display a summary of the selected files.

6. Download the resulting CSV file(s) containing the expression matrices.

2. **Download by Project**

   - **Purpose**: Download specific to individual BioProjects associated with this study.
   - **Steps**:
     1. Navigate to **Downloading Data > Download by Project**.
     2. View the table listing available BioProjects and their details.
     3. Select one or multiple rows corresponding to the desired projects.
     4. Click **Download Selected Project Data** on the right side to download the data for the selected projects.

3. **Latest Annotation**

   - **Purpose**: Download the latest genome annotation file (GFF3) created in this study.
   - **Steps**:
     1. Navigate to **Download > Latest Annotation**.
     2. Click the download link to retrieve the GFF3 file containing the latest *Artemisia annua* annotations.

4. **Latest Transcriptome Assembly**

   - **Purpose**: Download the latest transcriptome assembly file (FASTA) created in this study.
   - **Steps**:
     1. Navigate to **Download > Latest Transcriptome Assembly**.
     2. Click the download link to retrieve the FASTA file containing the latest *Artemisia annua* transcriptome assembly.

---

#### About the Database

Learn about the project, team, and future plans.

---

#### Tips and Troubleshooting

- **Contact**: Reach out via GitHub (https://github.com/ayattaheri) or email for support.

---

This tutorial provides a foundation for users to explore the Artemisia Database. For advanced usage or specific research questions, refer to the source code or contact the developer. Happy exploring!

---
